# Biochemical and Structural Characterization of Fapy•dG Replication by Human DNA Polymerase β

**DOI:** 10.1101/2024.01.15.575758

**Authors:** Shijun Gao, Peyton N. Oden, Benjamin J. Ryan, Haozhe Yang, Bret D. Freudenthal, Marc M. Greenberg

## Abstract

N6-(2-deoxy-α,β-D-erythro-pentofuranosyl)-2,6-diamino-4-hydroxy-5-formamido-pyrimidine (Fapy•dG) is formed from a common intermediate and in comparable amounts to the well-studied mutagenic DNA lesion 8-oxo-7,8-dihydro-2’-deoxyguanosine (8-OxodGuo). Fapy•dG preferentially gives rise to G → T transversions and G → A transitions. However, the molecular basis by which Fapy•dG is processed by DNA polymerases during this mutagenic process remains poorly understood. To address this we investigated how DNA polymerase β (Pol β), a model mammalian polymerase, bypasses a templating Fapy•dG, inserts Fapy•dGTP, and extends from Fapy•dG at the primer terminus. When Fapy•dG is present in the template, Pol β incorporates TMP less efficiently than either dCMP or dAMP. Kinetic analysis revealed that Fapy•dGTP is a poor substrate but is incorporated ∼3-times more efficiently opposite dA than dC. Extension from Fapy•dG at the 3’-terminus of a nascent primer is inefficient due to the primer terminus being poorly positioned for catalysis. Together these data indicate that mutagenic bypass of Fapy•dG is likely to be the source of the mutagenic effects of the lesion and not Fapy•dGTP. These experiments increase our understanding of the promutagenic effects of Fapy•dG.

## INTRODUCTION

2’-Deoxyguanosine (dG) oxidation has been extensively studied due to its relative ease of oxidation compared to the other native 2’-deoxynucleotides, as well as the general biological importance of DNA damage (1,2). Issues examined include product (lesion) identification, their formation levels, mutagenic properties and characterization of the interactions of lesions with DNA damage response enzymes (3,4). Of the lesions formed from dG oxidation, 8-oxo-7,8-dihydro-2’-deoxyguanosine (8-OxodGuo) has garnered the greatest attention (Scheme 1) (1,5–7). In addition to promoting mutagenesis, this lesion is considered as a biomarker and may play an epigenetic role in regulating gene expression (8–12). N6-(2-deoxy-α,β-D-erythro-pentofuranosyl)-2,6-diamino-4-hydroxy-5-formamido-pyrimidine (Fapy•dG) is a DNA lesion derived from a common precursor (**1**) to 8-OxodGuo (Scheme 1) (13). Moreover, it is produced in amounts comparable to those of 8-OxodGuo in some cell lines and under anoxic conditions *in vitro* (14–19). The cleavage of the 5-membered ring of the purine introduces unusual structural properties. Release of the lone pair electrons on the glycosidic nitrogen from the aromatic systems endows Fapy•dG with the unusual ability to isomerize between α- and β-anomers via an acyclic intermediate (Scheme 1) (20). In addition, the formamide group is free to rotate, as is the entire pyrimidine ring about the carbon-nitrogen bond. The latter can result in flipping the orientation of the Watson-Crick face and positioning the formamide group in either the major or minor groove. The ability of Fapy•dG to adopt multiple stereoisomers (conformational and configurational) increases the possible ways in which a polymerase and other repair proteins can interact with it. This could result in diverse outcomes compared to more rigid lesions, including 8-OxodGuo. However, there is a significant knowledge gap concerning nucleic acid structure containing Fapy•dG and its recognition by enzymes involved in the DNA damage response. A recently reported improved method for synthesizing oligonucleotides containing Fapy•dG has made such studies on this lesion more tractable (21,22). Herein, we examine biochemical and structural aspects of DNA replication involving Fapy•dG and Fapy•dGTP using human DNA polymerase β (Pol β).

**Scheme 1.**
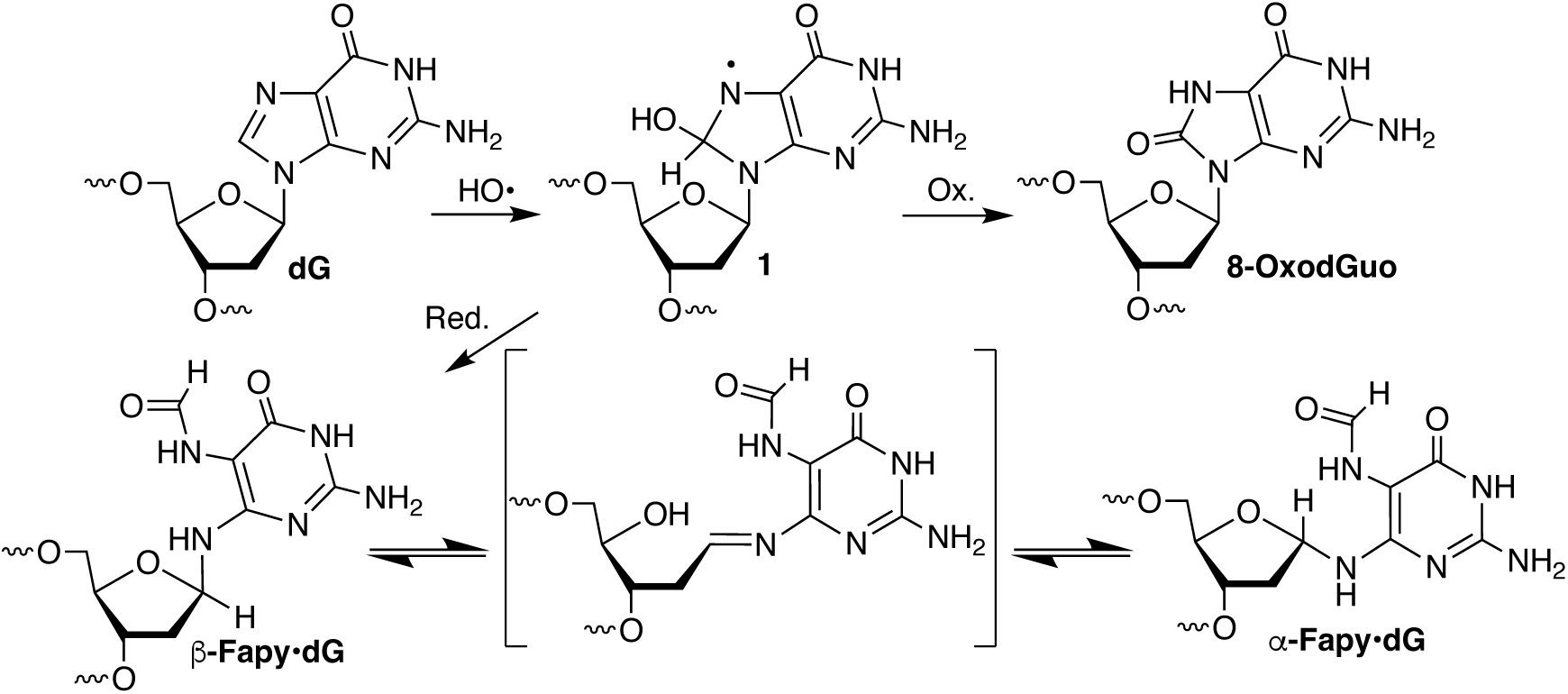
Fapy•dG and 8-OxodGuo formation from a common intermediate.

Investigations using single-stranded vectors revealed that Fapy•dG replication in mammalian cells (HEK 293T, COS-7), including at mutation hotspots within the p53 gene, is more mutagenic than 8-OxodGuo (23–25). There are also significant differences in the distribution of promutagenic events when Fapy•dG and 8-OxodGuo are bypassed. G → T transversion point mutations are the predominant event when 8-OxodGuo is bypassed (12,25,26). Fapy•dG bypass yields a more diverse spectrum of promutagenic products that includes G → A transitions in addition to G → T transversions, and even G → C transversions and single-nucleotide deletions (23). Translesion incorporation of dCMP and dAMP opposite Fapy•dG by Pol β has been characterized kinetically and structurally (27). X-ray structure analysis reveals that the α-/ β-Fapy•dG equilibrium responds to the opposing nucleotide when a nonhydrolyzable dNTP analogue is part of a ternary complex comprised of Pol β and a primer-template DNA substrate. The adjustment of the Fapy•dG configuration to the local environment is a distinctive structural property that cannot be revealed by studies on configurationally stable analogues (28). Comparable information regarding TMP incorporation opposite Fapy•dG that is relevant to the formation of G → A transitions in cells is unavailable. Although kinetic information was reported for extension past Fapy•dG by the bacterial polymerase Klenow exo^−^, this current study addresses a gap in our knowledge regarding bypass by mammalian DNA polymerases (29).

Promutagenic base pairs are also produced as a result of oxidation of the dNTP pool and subsequent damaged nucleotide monophosphate incorporation into DNA by polymerases. Digested food is another source of damaged dNTPs that find their way into cellular DNA (30). Pol β incorporates 8-OxodGuoMP opposite dA ∼40-times more efficiently than opposite dC, promoting genomic mutations (31). Structural and kinetic analysis of extension from the inserted 8-OxodGMP opposite dA or dC by Pol β revealed this process could be detrimental to a cell by leading to aborted DNA synthesis or mutagenesis because extension from 8-OxodGMP opposite dA is favored in comparison to 8-OxodGMP opposite dC (31,32). MutT (bacteria) and MTH1 (mammalian cells) protect the genome by sanitizing the dNTP pool of damaged dNTPs such as 8-OxodGuoTP via hydrolysis of a pyrophosphate bond. The importance of MTH1 sanitizing the pool to protect genome stability is underscored by promising anticancer agents that alter the activity of MTH1 (33–35). The ability of MTH1 to hydrolyze Fapy•dGTP is unknown, but the triphosphate is a poor substrate for MutT (36). The specificity constant for Fapy•dGTP hydrolysis by MutT is 4-orders of magnitude slower than that of 8-OxodGuoTP and 100-fold less than that of dGTP. The weak activity of MutT on Fapy•dGTP likely increases the levels in the nucleotide pool and potential for a DNA polymerase to insert it during DNA synthesis. Klenow exo^−^ incorporates Fapy•dGTP opposite dC 25-times more rapidly than when dA is present in a template but the enzyme utilizes Fapy•dGTP 1,000-fold less efficiently than dGTP (36). Although mammalian polymerase data were unavailable with Fapy•dGTP as a substrate, Pol β incorporates a C-nucleotide analogue (β-C-Fapy•dGTP) opposite dC 82-times more efficiently than opposite dA (37). However, the specificity constant for β-C-Fapy•dGTP is more than 5,000-times lower than dGTP. The use of Fapy•dGTP as a substrate by Pol β and the subsequent extension of a primer containing Fapy•dG at the 3’-teminus is addressed below.

## MATERIAL AND METHODS

### General materials and methods

[γ-^32^P]-ATP was from Perkin Elmer Health Sciences Inc. (Shelton, CT). dNTPs were purchased from New England Biolabs (Beverly, MA). Oligonucleotides were either synthesized on an ABI 394 oligonucleotide synthesizer at JHU or obtained from Integrated DNA Technologies (Coralville, IA). Fapy·dG-containing oligonucleotides were synthesized as described previously (21), purified by 20% denaturing polyacrylamide gel electrophoresis and characterized by MALDI-TOF MS (Figure S2 and S3). PC Spacer phosphoramidite for 5’-phosphorylation, 5’-cyanoethyl phosphoramidites for thymidine (T), N,N-dimethylformamidine-2’-deoxyguanosine (dG^dmf^), as well as other reagents required for oligonucleotide synthesis were purchased from GLEN Research (Sterling, VA). N-acetyl 2’-deoxycytidine 5’-cyanoethyl phosphoramidite (dC^Ac^) and Universal UnyLinker™ Support were purchased from Chemgenes (Wilmington, MA). 5’-cyanoethyl phosphoramidite of N-Pac-2’-deoxyadenosine was prepared as previously described (22). Disodium 2-carbamoyl-2-cyanoethylene-1,1-dithiolate trihydrate for methyl deprotection was prepared as previously described (38).

### Expression and Purification of Polymerase β

BL21-CodonPlus(DE3)-RP Escherichia coli cells were used to overexpress wild-type human DNA Pol β in a pET-30a vector. Protein expression was induced when the OD600 reached 0.6 with 0.1 mM isopropyl-β-D-thiogalactopyranoside (IPTG). Temperature was reduced to 18 °C and cells were harvested the next day. As previously described, protein purification was performed (39). *E. coli* Cell lysate was run over a HiTrap Heparin HP, Resource S, and HiPrep 16/60 Sephacryl S-200 HR column. Fractions containing Pol β were ran on an SDS page gel to confirm purity. Pure pol β was concentrated and aliquoted. Aliquots for crystallization were stored in 20 mM BisTris propane, pH 7.0 at −80 °C. Aliquots for kinetic experiments were stored in 50 mM Hepes pH 7.5, and 150 mM NaCl at −80 °C.

### X-Ray Crystallography DNA Sequences

Oligonucleotides were resuspended in 10 mM Tris-HCl, pH 7.4 and 1 mM EDTA. Concentration of oligonucleotides was measured using ultraviolet absorbance at 260 nm. Crystallization studies with Fapy•dG in the template strand, opposite the primer terminus, used the following DNA sequence to generate 16-mer gapped DNA substrates: 5’ – CCGACC<u>F</u>CGCATCAGC – 3’, where <u>F</u> represents Fapy•dG. The downstream primer sequence used for all studies was: 5’ – pGTCGG – 3’. The upstream primer used was: 5’ – GCTGATGCG<u>X</u> – 3’, where <u>X</u> represents dC, dA, or T. The following DNA template sequence was used for crystallization studies with Fapy•dG at the primer terminus: 5’ – CCGACG<u>X</u>CGCATCAGC – 3’. X represents dC, dA or T. The upstream primer used was: 5’ – GCTGATGCG<u>F</u> – 3’ (21). Further, 16-mer DNA substrates were also used in crystallization studies with Fapy•dG as the templating base. The 16-mer DNA sequence is as follows: 5’ – CCGAC<u>F</u>TCGCATCAGC – 3’ (21). The upstream primer used in the study was: 5’ – GCTGATGCGA – 3’. Host-Guest complex crystallization studies used the following 16-mer: 5’ – CCGACGGCGCAT<u>F</u>AGC – 3’ (27). The upstream primer used was: 5’ – GCT<u>T</u>ATGCGC – 3’. The gapped DNA substrates were made by annealing three oligonucleotides: template, downstream, and upstream. The oligonucleotides were incubated in a solution at 95 °C for 5 minutes, 65 °C for 30 minutes, and cooled to 4 °C until use in a PCR thermocycler.

### Kinetics of Pol β mediated incorporation of Fapy•dGMP opposite dC/dA/T

To prepare the gapped DNA complex **2a-c**, the 5’-^32^P-labeled primer was mixed with 1.2 equivalents of template and 1.2 equivalents 5’-phosphorylated flanking strand in 1× PBS buffer (137 mM NaCl, 2.7 mM KCl, 10 mM Na_2_HPO_4_, and 1.8 mM KH_2_PO_4_, pH 7.5). The mixture was heated to 60 °C or 5 min, followed by a slow cooling to room temperature. The gapped DNA complex was mixed with DNA polymerase β (Pol β) to create 2× DNA-enzyme solution (DNA 200 nM, Pol β 40 nM) in 1× reaction buffer (50 mM Tris-HCl pH 7.5, 100 mM KCl, 1 mM DTT, 5 mM MgCl_2_, 0.1 mg/mL BSA, 10% glycerol). The DNA-enzyme mixture was incubated on ice for 5 min. Next, an equal volume of 2× Fapy·dGTP solution in 1× reaction buffer was added to initiate the reaction. Final concentration: DNA 100 nM, Pol β 20 nM. If the desired reaction time is shorter than 3 min, the DNA-enzyme mixture was incubated at 37 °C for at least 1 min prior to the addition of 2× Fapy·dGTP solution. The concentration of Fapy·dGTP and reaction time were chosen to ensure that the reaction did not proceed to more than 30% completion. To quench the reaction, 2 volumes of the quenching solution (90% formamide, 50 mM EDTA, 0.05% xylene cyanol and 0.05% bromophenol blue) was added. Samples were denatured at 95 °C for 5 min before loading onto a 20% denaturing polyacrylamide gel. Velocities were determined by the equation [100I_1_/(I_0_ + 0.5I_1_)]t, where I_1_ is the signal of product, I_0_ is the signal of starting material, t is the reaction time (40). In each experiment, reactions were carried out in triplicate. The Fapy·dGTP concentrations and velocities were fitted to the Michaelis–Menten equation to determine *k_cat_* and *K_M_*.

The Fapy·dGTP concentrations and reaction time are described below:

For **2a**, [Fapy·dGTP] 30, 60, 120, 200, 500, 1000 μM, 10 min for 30, 60, 120 μM and 2 min for 200, 500, 1000 μM;
For **2b**, [Fapy·dGTP] 30, 60, 120, 200, 500, 1000 μM, 15 min for 30 μM, 5 min for 60 μM, 2 min for 120 μM, 1 min for 200 μM, 40 s for 500 μM and 1000 μM;
For **2c**, [Fapy·dGTP] 30, 60, 120, 200, 500, 1000 μM, 10 min for 30 μM, 6 min for 60 μM, 3 min for 120 μM, 1 min for 200 μM, 40 s for 500 μM and 1000 μM.

### Pol β mediated time-course extension of Fapy·dG opposite dC/dA/T

To prepare the gapped DNA complex **4a-c**, the 5’-^32^P-labeled primer was mixed with 1.2 equivalents of template and 1.2 equivalents 5’-phosphorylated flanking strand in 1× PBS buffer. The mixture was heated to 60 °C for 5 min, followed by a slow cooling to room temperature. The ternary DNA complex was mixed with DNA polymerase β to create 2× DNA-enzyme solution (DNA 200 nM, Pol β 40 nM) in 1× reaction buffer. The DNA-enzyme mixture was incubated on ice for 5 min. Next, an equal volume of 2× dGTP solution in 1× reaction buffer was added to initiate the reaction. Final concentration: DNA 100 nM, Pol β 20 nM. At each selected time point, an aliquot of the reaction was added to 2 volumes of the quenching solution (90% formamide, 50 mM EDTA, 0.05% xylene cyanol and 0.05% bromophenol blue). Samples were denatured at 95 °C for 5 min before loading onto a 20% denaturing polyacrylamide gel.

The dGTP concentrations and time points are described below:

Extension of **4a**, [dGTP] 1 μM, 5 μM, 10 μM. Time points: 3 min, 10 min, 30 min;
Extension of **4b**, [dGTP] 100 μM, 300 μM, 1000 μM. Time points: 3 min, 10 min, 30 min;
Extension of **4c**, [dGTP] 100 μM, 300 μM, 1000 μM. Time points: 3 min, 10 min, 30 min.

### Kinetics of Pol β mediated extension of Fapy·dG opposite dC

The experiment was done in the same manner as described for incorporation of Fapy·dGTP using ternary DNA complex **4a**. Final concentration: DNA 100 nM, Pol β 20 nM.

dGTP concentrations and reaction time are described below:

[dGTP] 2, 5, 10, 20, 50, 100 μM, 3 min for 2 μM and 5 μM, 1 min for 10 μM and 20 μM, 30 s for 50 μM and 100 μM.

### Kinetics of Pol β mediated incorporation of TTP opposite Fapy·dG

The experiment was done in the same manner as described for incorporation of Fapy·dGTP using ternary DNA complex **6**. Final concentration: DNA 100 nM, Pol β 3 nM

The TTP concentrations and reaction time are described below:

[TTP] 50, 150, 300, 500, 750, 1000 μM, 2 h for all concentrations.

### Kinetics of Pol β mediated extension of dC/dA/T opposite Fapy·dG

The experiment was done in the same manner as described for incorporation of Fapy·dGTP using ternary DNA complex **7a-c**. Final concentration: DNA 100 nM, Pol β 3 nM.

The dGTP concentrations and reaction time are described below:

Extension of **7a**, [dGTP] 2, 5, 10, 20, 50, 100 μM, 3 min for 2 μM and 5 μM, 1 min for 10 μM and 20 μM, 40 s for 50 μM and 100 μM;
Extension of **7b**, [dGTP] 10, 20, 50, 100, 200, 500 μM, 10 min for all concentrations;
Extension of **7c**, [dGTP] 10, 20, 50, 100, 200, 500 μM, 3 min for 10 μM and 20 μM, 1 min for 50 μM and 100 μM, 40 s for 200 μM and 500 μM.

### X-Ray Crystallography

Vapor diffusion was used for protein crystallization. The crystallization solution was made with 14-22% PEG 3350, 0.05 M Imidazole pH 8.0, and 0.35 Sodium Acetate. Annealed DNA at a concentration of 0.5 mM was incubated with Pol β (7-12 mg/mL) for 15 minutes before the addition of the crystallization solution, mentioned above. Crystals were soaked in a cryoprotectant comprised of 20% ethylene glycol and 80% crystallization solution before mounting and shooting. A soak consisting of a nucleotide included a catalytic metal (MgCl_2_ or MnCl_2_) and the preferred nucleotide at a concentration of 10 mM and 1 mM, respectively. Soaks consisting of a nucleotide incubated for at least 1 hour to allow for nucleotide insertion. Crystals were mounted and shot with a Dectris Pilatus3R 200 K detector system. Data was collected on a Rigaku MicroMax-007 HF rotating anode diffractometer at 100 K. Protein models were produced with Phaser-MR (simple one-component interface) using PDB files of Pol β binary and ternary complexes (PDB ID code: 2FMS, 3ISB, 7S9O). PHENIX and Coot were used to refine and build the models (41,42). Figures in the publication were produced using PyMOL (Schrödinger).

## RESULTS

### Fapy•dGMP incorporation and subsequent oligonucleotide extension

Fapy•dGTP was synthesized as previously described (36). Utilization of Fapy•dGTP by Pol β as a substrate was examined using single-nucleotide gapped complexes (Table 1, **2a-c**) (41). The gapped complexes contained dC, dA or T in the template strand at the position opposite to where Fapy•dGMP would be inserted (“N” in Table 1, **2a-c**). The corresponding complex containing dG was not quantitively examined, because significant levels of G → C transversions are not observed upon Fapy•dG replication in cells. The specificity constant for lesion incorporation opposite dA was greatest, ∼3-fold greater than opposite dC (Table 1). Although the specificity constant for incorporation opposite T (**2c**) was almost as high as when dA was in the template, the *K*_M_ was extremely high (∼ 1 mM). The difference in specificity constants between when dA (**2b**) or dC (**2a**) was present in the template was due to *V*_max_, as the *K*_M_’s were within experimental error of one another. The preference for incorporating Fapy•dGMP opposite dA over dC by Pol β is also observed when 8-OxodGuoTP is the substrate (31,42). However, Pol β was more discriminating, preferring to incorporate 8-OxodGuoMP opposite dA as much as 40-times more efficiently than when dC was present in the template. Moreover, 8-OxodGuoTP is a much better substrate for Pol β than is Fapy•dGTP. The specificity constant for 8-OxodGuoTP when dA is in the template is ∼1,000-times higher than for Fapy•dGTP (Table 1) (31). Although *V*_max_ and *K*_M_ values are not reported, pre-steady state experiments indicate that 8-OxodGTP is bound more strongly by Pol β, particularly when dA is the opposing nucleotide in the template (42). In addition, the polymerization rate constant for introducing 8-OxodGMP opposite dA is only ∼6-fold lower than the respective rate constant for incorporation of dGMP opposite dC. The relatively inefficient specificity constant for forming product (**3a-c**) may explain why, despite significant effort, we were unable to obtain structural data when soaking Pol β-DNA cocrystals with Fapy•dGTP.

**Table 1.** Fapy•dGMP incorporation by Pol β.^a^.

| N | $k_{\text{cat}}$ ($\times 10^{-2} \text{ s}^{-1}$) | $K_{\text{M}}$ ($\mu\text{M}$) | $k_{\text{cat}} / K_{\text{M}}$<br>($\times 10^{-5} \mu\text{M}^{-1} \text{ s}^{-1}$) |
| --- | --- | --- | --- |
| C<sup>b</sup> | 1.2 ± 0.2 | 346.8 ± 29.8 | 3.5 ± 0.6 |
| A<sup>b</sup> | 3.1 ± 0.1 | 345.4 ± 14.1 | 9.0 ± 0.4 |
| T<sup>c</sup> | 8.2 ± 1.1 | 1015.3 ± 263.8 | 8.1 ± 2.4 |

<sup>a</sup>[DNA] = 100 nM, [Pol $\beta$ ] = 20 nM. <sup>b</sup>Data are the ave. ± std. dev. of 3 experiments. Each experiment is comprised of 3 replicates. <sup>c</sup>Data are the ave. ± std. dev. of 1 experiment containing 3 replicates.

To complete lesion incorporation, a nascent primer must be extended. We explored this step using ternary DNA complexes containing Fapy•dG in the primer strand at the 3’-terminus opposite dC, dA or T (Scheme 2, **4a-c**). Significant strand extension (Scheme 2, **5a-c**; [**4a-c**] = 100 nM, [Pol β] = 20 nM) in the presence of physiologically relevant dGTP concentrations (≤ 10 μM) (43) was only observed within 3 min when Fapy•dG was opposite dC (**4a**, Figure S1). The specificity constant ((3.9 ± 0.6) × 10^−3^ μM^−1^s^−1^) for this process is ∼80-fold less efficient than when the gapped complex is comprised entirely of native nucleotides (41). However, it is significantly more efficient than when Fapy•dG is opposite dA or T. When Fapy•dG is opposite T (**4c**), significant product (**5c**) is observed only at higher, physiologically irrelevant dGTP concentrations (100 – 1,000 μM) and longer reaction times (3 – 30 min). Extension was even less efficient when Fapy•dG was opposite dA (**4b**).

**Scheme 2.**
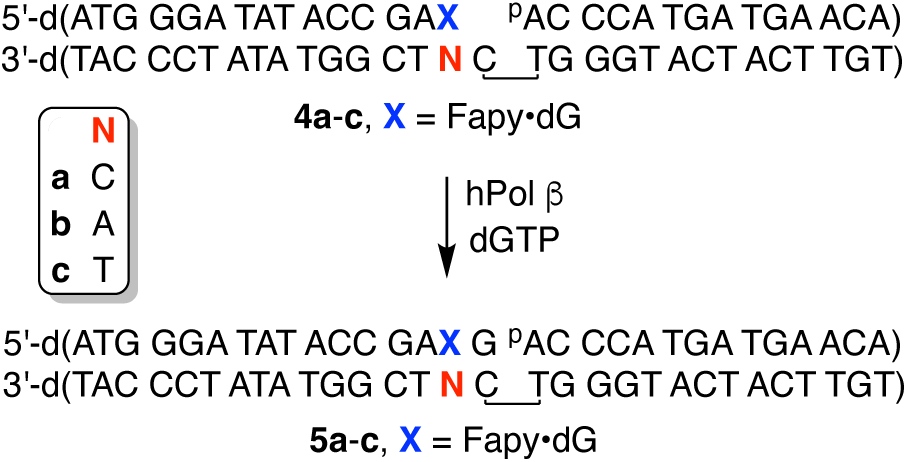
Single nucleotide extension of DNA containing 3’-Fapy•dG.

The Pol β extension experiments were corroborated by X-ray crystallography studies in which we characterized Fapy•dG at the primer 3’-terminal position opposite dC, dA, and T. Initial characterization of a binary Fapy•dG-dC was accomplished via crystallization of Pol β and Fapy•dG-dC containing DNA. The resulting crystal diffracted to 1.69 Å and was solved in space group P2_1_ with Pol β in the open conformation (Table S1, Figure 1a). Fapy•dG forms a Watson-Crick base pair with dC consisting of typical hydrogen bond distances and we only observed the β-Fapy•dG anomer (Figure 1b). The pyrimidine ring adopts a conformation such that the formamide group is positioned in the major groove of DNA and is coordinated by a single ordered water molecule (Figure 1b). The Fapy•dG-dC base pair is in a stable conformation, as indicated by clear electron density and similar average B-factors of 25.2 Å^2^ for Fapy•dG and 26.7 Å^2^ for dC. Overall, the active site is in position to allow binding of the incoming nucleotide.

**Figure 1.**
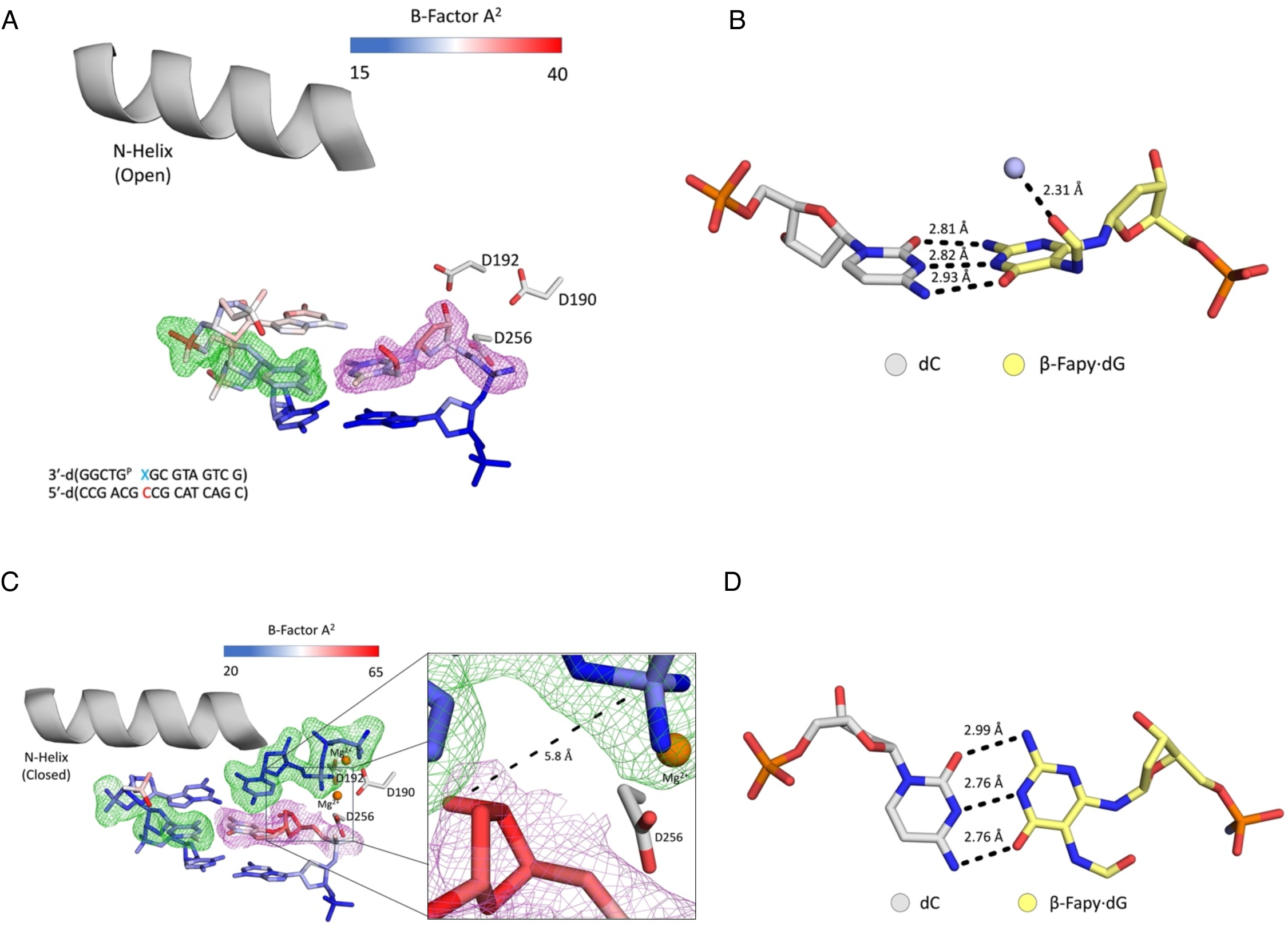
dC opposite Fapy•dG at the primer terminus. **(A)** Active site of the binary Pol β complex. A legend for the B-factors is shown in a blue-white-red gradient. **(B)** β-Fapy•dG (yellow sticks) base paired with dC (gray sticks) with a water molecule shown as a light blue sphere. **(C)** Ternary Pol β complex with an incoming dCTP. The inset shows a focused view of the 3′-OH and α-PO_4_. **(D)** β-Fapy•dG (yellow sticks) base paired with dC (gray sticks) from the ternary complex. Magnesium ions are colored orange. Omit maps for are shown as green mesh contoured to 2σ. Polder maps are shown as purple mesh contoured to 3σ for A and 2σ for C. Key distances are shown as dashes and labels.

Further characterization of extension from Fapy•dG-dC was achieved by solving a ternary (DNA:dNTP:Pol β) structure of primer terminal Fapy•dG-dC with an incoming dCTP. This structure was solved by soaking a binary crystal in a cryo-solution containing 1 mM dCTP and 10 mM MgCl_2_. The resulting crystal diffracted to 2.01 Å in space group P2_1_. The Fapy•dG-dC primer terminal base pair and the incoming dCTP are accommodated in the active site with the dCTP base pairing to the templating dG. Pol β is in a closed conformation and only β-Fapy•dG is observed (Figure 1c). Fapy•dG base pairs to dC with its Watson-Crick face with standard hydrogen bond distances (Figure 1d). Similar to the binary structure, the formamide group is positioned in the major groove of DNA. In this structure, Fapy•dG has an increased B-factor_avg_ (48.4 Å^2^) compared to the entire duplex DNA (40.6 Å^2^). Furthermore, the B-Factor of Fapy•dG’s 3’-OH is substantially higher than the entire residue at 57.9 Å^2^ (Figure 1c). The 3’-OH sits in the minor groove, 5.8 Å away from the α-PO_4_ of the incoming dCTP, which is not in a catalytically competent position to coordinate the catalytic metal (Figure 1c). While we observe nucleotide binding, we do not observe insertion of the dCMP likely due to the position of the 3’-OH. This is in accordance with the inefficient extension reflected by the ∼80-fold reduction in specificity constant of (**4a**) compared to when native dG is present.

To characterize extension after promutagenic incorporation of Fapy•dG opposite dA (Fapy•dG-dA), Pol β:DNA crystals containing primer terminal Fapy•dG-dA were obtained. We solved a binary structure at 1.69 Å in space group P2_1_ with Pol β in the open conformation (Table S1 and Figure 2a). The omit map shows that Fapy•dG is very dynamic with poor density, which contrasts with Fapy•dG-dC (Figure 1a and 2a). The best model indicates the Watson-Crick face of Fapy•dG engages in a single hydrogen bond with the opposing dA (Figure 2b). Due to lack of a single stereoisomer in the density of the omit map for Fapy•dG, its configuration cannot be determined and is modeled as the β-anomer. The average B-factor of Fapy•dG at the primer terminus is 31.8 Å^2^, while the B-factor of the 3’-OH of Fapy•dG is 38.3 Å^2^. These B-factors are substantially higher compared to the rest of the DNA (B-factor = 22.3 Å^2^). Of note, the dA opposite Fapy•dG shifts 1.2 Å to accommodate the lesion and mispairing, compared to a matched non-damaged primer terminus (Figure S4a and S4b).

**Figure 2.**
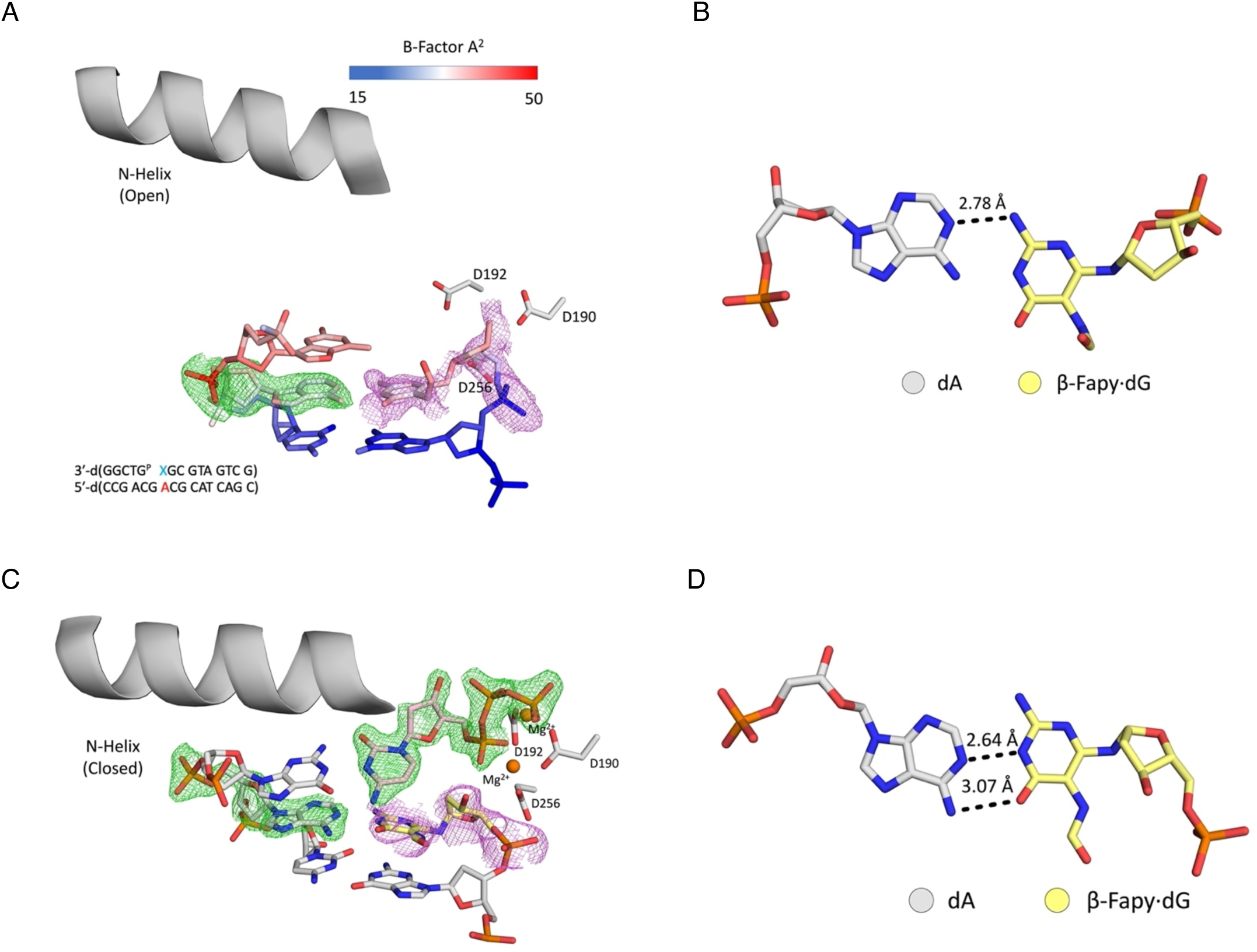
Primer terminal Fapy•dG base-paired with dA. **(A)** Active site of the binary Pol β complex. A legend for the B-factors is shown in a blue-white-red gradient. **(B)** β-Fapy•dG (yellow sticks) base paired with dA (gray sticks). **(C)** Ternary Pol β complex with an incoming dCTP. **(D)** β-Fapy•dG (yellow sticks) base paired with dA (gray sticks) from the ternary complex. Magnesium ions are colored orange. Omit maps are shown as green mesh contoured to 2.5σ for A and 1.5σ for C. Polder maps are shown in purple mesh contoured to 2.2σ for A and 2σ for C. Key distances are shown as dashes and labeled.

To further characterize the extension of Fapy•dG-dA, we solved a ternary structure with an incoming dCTP by soaking a binary crystal in a cryo-solution containing 1 mM dCTP and 10 mM MgCl_2_. The resulting crystal diffracted to 1.77 Å in space group P2_1_ with Pol β in the closed conformation. The Fapy•dG-dA primer terminal base pair, as well as the incoming dCTP, is accommodated in the active site with the dCTP base pairing to the templating dG. (Table S1, Figure 2c). The density for Fapy•dG at the primer terminus is poor, suggesting it is dynamic (Figure 2c). Due to lack of a clear single stereoisomer in the density for Fapy•dG, the configuration of the lesion cannot be determined and is modeled as the β-anomer. In contrast to when dCTP is absent, Fapy•dG interacts with dA via hydrogen bonds between the N1 of dA and the N1 of Fapy•dG, as well as the N6 of dA and the O6 of Fapy•dG (Figure 2d). The primer terminal 3’-OH of Fapy·dG is 3.6 Å from the α-phosphate of the incoming dCTP, but is not coordinated by the catalytic metal (4.2 Å away from the metal). This differs from an undamaged nucleotide with respect to both distances (Figure S5a). Comparison of Fapy•dG to a non-damaged nucleotide reveals extensive conformational changes at the primer terminus, including a 3.2 Å shift of the 3’-OH into the major groove and the absence of catalytic metal coordination (Figure S5b). The complex would need to undergo adjustements in the active site to allow for catalysis. This structural snapshot would require additional active site adjustments to allow catalysis and is consistent with the inefficient extension we observed in kinetic studies of **4b**.

Given one of the mutagenic signatures for Fapy•dG is G → A transitions, we set out to understand the structural dynamics of the extension of primer terminal Fapy•dG-T base pair. To this end we solved a binary structure at 2.09 Å in space group P2_1_ with Pol β in the open conformation (Figure 3a). Fapy•dG uses its Watson-Crick face to form a single hydrogen bond between the O4 of T and the N2 of Fapy•dG (Figure 3b). The Fapy•dG epimerization state cannot be determined due to insufficient density and is modeled as the β-anomer. Rotation about the glycosidic nitrogen-pyrimidine carbon bond places the formamide group out of plane and in the DNA minor groove. Primer terminal Fapy•dG-T results in a very dynamic active site indicated by the poor density for Fapy•dG, (Figure 3a), which is consistent with the observed inefficient extension of a corresponding ternary complex (Scheme 2, 4c). We were unable to obtain a ternary structure with primer terminal Fapy•dG-T likely due to instability of the primer terminus preventing binding of the incoming dCTP.

**Figure 3.**
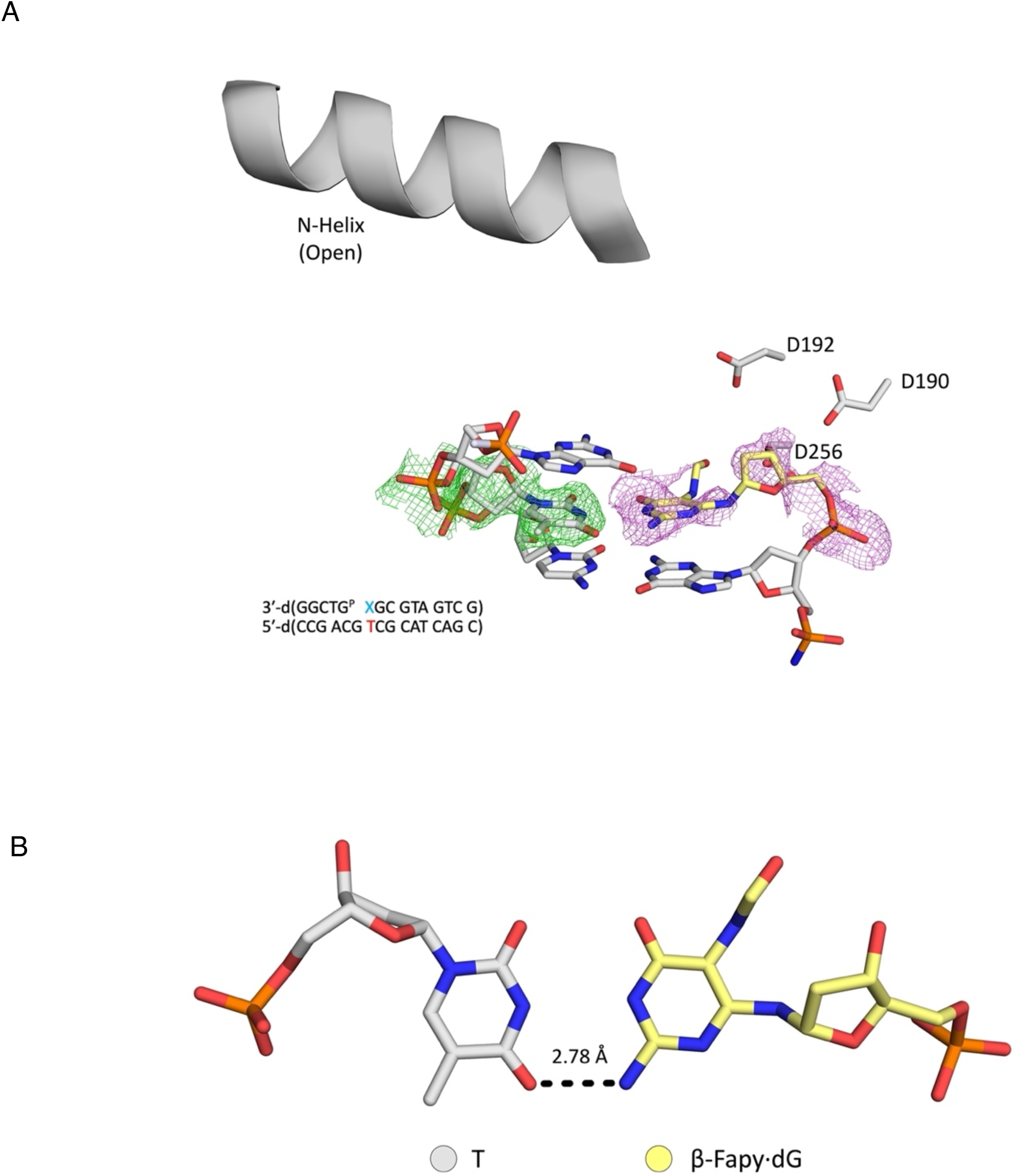
Primer terminal Fapy•dG base-paired with T. **(A)** Binary Pol β complex with with Fapy•dG at the primer terminus, opposite T. **(B)** β-Fapy•dG (yellow sticks) base paired with T (gray sticks). Omit map is shown as green mesh contoured to 2σ. Polder map is shown as purple mesh contoured to 2σ.

### TMP incorporation opposite Fapy•dG

The relative frequency of G → A transitions compared to G → T transversions when Fapy•dG is bypassed in HEK 293T cells ranged from approximately 1/6^th^ to roughly equal within the three sequences examined (23). Kinetic and structural studies on dCMP and dAMP incorporation opposite Fapy•dG have provided insight into G → T transversions (27). Here, we determined the kinetic constants (*K*_M_ = 601 ± 115 μM, *k*_cat_ = (1.5 ± 0.2) × 10^−3^ s^−1^, *k*_cat_ / *K*_M_ = (2.5 ± 0.6) × 10^−6^ μM^−1^s^−1^) for TMP incorporation opposite Fapy•dG in **6** (Scheme 3). The specificity constant was more than 20,000-times less efficient than dCMP incorporation and ∼15-times less than for dAMP incorporation (27). A greater *K*_M_, which was ∼20-fold and 600-fold higher for TTP than that of dATP and dCTP, respectively, were greater contributors to distinguishing between specificity constants than were *k*_cat_ values.

**Scheme 3.**
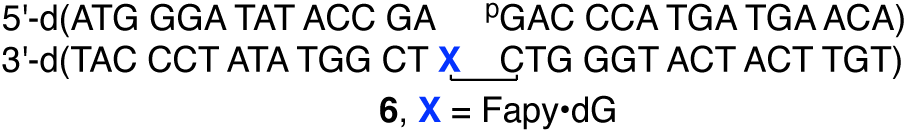
Ternary DNA complex for examining TTP incorporation opposite Fapy•dG.

A structural understanding of TTP incorporation opposite Fapy•dG was identified via X-ray crystallography using crystals of Pol β in complex with DNA containing Fapy•dG in the templating position across from a nonhydrolyzable TTP analog (dTpCpp). Binary crystals were soaked in a cryo-solution containing 1 mM dTpCpp and 10 mM MnCl_2_. MnCl_2_ was used to overcome the poor *K*_m_ for TTP binding as observed by TMP insertion kinetically and dTpCpp was utilized to prevent insertion. The resulting crystal diffracted to 2.10 Å and was solved in space group P2_1_. Pol β is in a partially closed conformation with the TTP analog bound (Figure 4a and Table S1). Fapy•dG presents a Watson-Crick face and only β-Fapy•dG is observed with the formamide group in the minor groove (Figure 4b). Although the incoming dTpCpp is bound in the active site, the dTpCpp is not within hydrogen bonding distance of the Fapy•dG. Strikingly, the B-factor of Fapy•dG (B-factor_avg_ = 61.5 Å^2^) is drastically higher than the B-factor of all the DNA residues in the structure (B-factor = 35.5 Å^2^), indicating increased dynamics. Importantly, the inability for dTpCpp to efficiently bind in a catalytically competent position provides a rationale for the extremely high *K*_m_ describing TMP incorporation opposite Fapy•dG.

**Figure 4.**
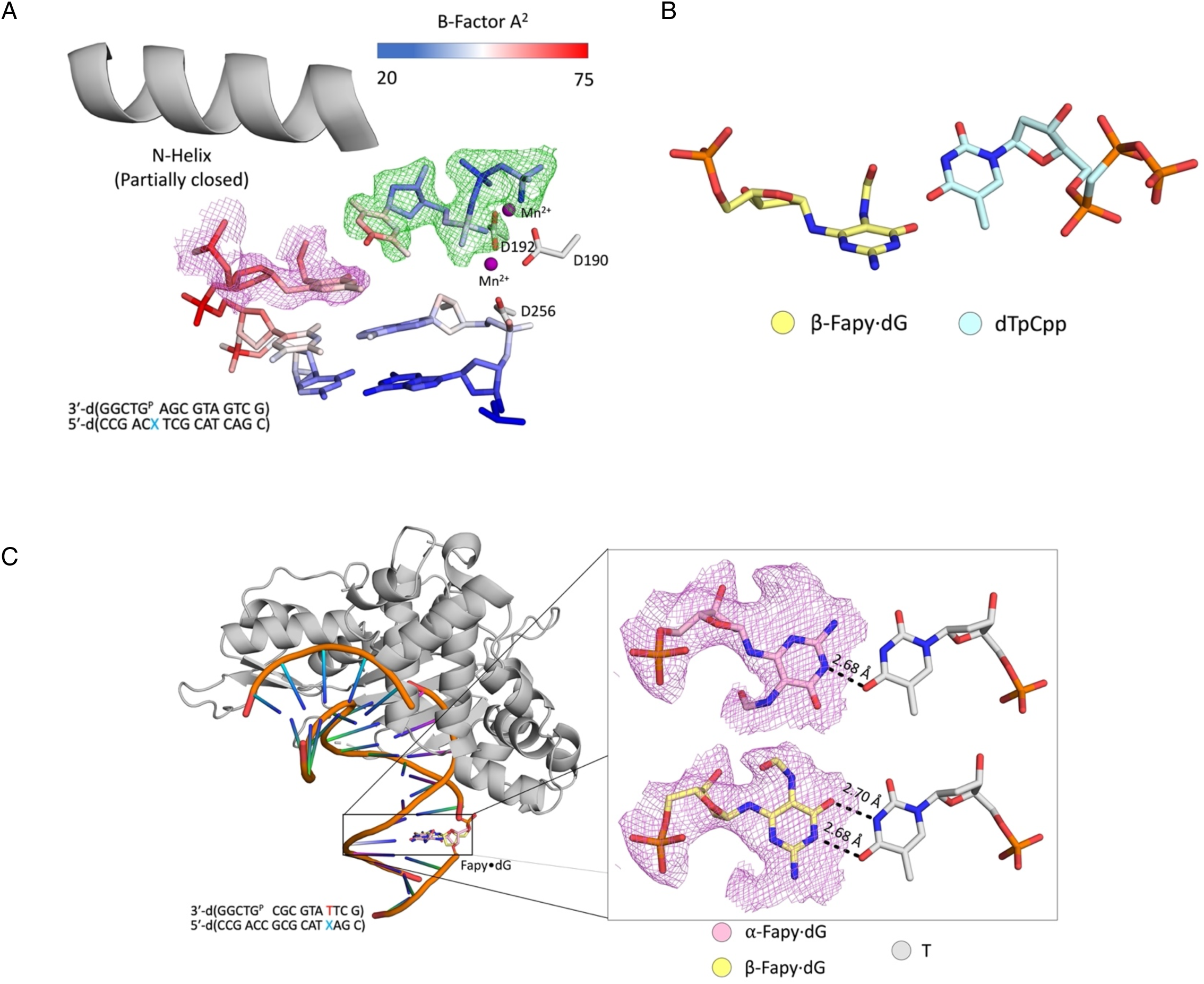
Structural effects when Fapy•dG is paired with thymidine. **(A)** Active site of the ternary Pol β:DNA complex. A legend for B-factors is indicated in the blue-white-red gradient. The omit map is shown as green mesh contoured to 2σ. The polder map (purple mesh) is contoured to 2σ. Manganese ions are colored purple. **(B)** Close up of the β-Fapy•dG (yellow sticks) incoming dTpCpp (cyan sticks) interaction. **(C)** Overall and inset view of Pol β (gray cartoon) host-guest complex. Inset shows α-Fapy•dG (pink sticks) and β-Fapy•dG (yellow sticks) base-paired with T (gray sticks). Manganese ions are colored purple. Polder maps are shown as purple mesh contoured to 2σ. Key distances are shown as dashes and labeled.

The Fapy•dG-T interaction was characterized further by analyzing the base pair combination in duplex DNA. To that end, we solved a Pol β:DNA crystal containing Fapy•dG-T host guest complex at 2.14 Å in space group P2_1_ (Figure 4c). Fapy•dG adopts α- and β-anomers when base paired with T in duplex DNA, with occupancy of β-Fapy•dG at 69% and α-Fapy•dG at 31%. The α-anomer forms one hydrogen bond between O4 of T and N1 of Fapy•dG (Figure 4c). In addition, the α-Fapy•dG pyrimidine ring adopts a conformation that places the formamide group in the major groove, which maintains the orientation of the Watson-Crick face presented by Fapy•dG in the same orientation as native dG. In contrast, the β-anomer interacts with T via two hydrogen bonds from O4 of T to N1 of Fapy•dG and N3 of T to O6 of Fapy•dG (Figure 4c). Furthermore, in the β-configuration the pyrimidine ring has rotated such that the formamide group is in the minor groove DNA and the Watson-Crick face is oriented in the opposite manner of native dG (Figure 4c). Reversing the orientation of the Watson-Crick face enables Fapy•dG to form two hydrogen bonds with the thymine ring. This structure indicates that once incorporated in the DNA, Fapy•dG-T is a relatively stable base pair.

### Primer extension following nucleotide incorporation opposite Fapy•dG

Pol β extension of the nascent strand containing dC opposite Fapy•dG (Table 2, **7a**), a nonmutagenic base pair, is ∼15-fold less efficient than when an undamaged DNA is acted on (Table 2) (41). The majority of the decrease in efficiency is due to a higher *K*_M_ for extension past Fapy•dG. Extension past a T-Fapy•dG base pair (**7c**) to form **8c** is ∼6-fold less efficient than when dC is opposite the lesion (**7a**). The slower specificity constant is attributable to a higher *K*_M_ for dGMP incorporation when T is opposite Fapy•dG; the corresponding *k*_cat_ is actually slightly higher. The primer-template containing a dA-Fapy•dG base pair (**7b**) is extended even less efficiently. Product (**8b**) is formed almost 60-times less efficiently than **8a**. Although the *K*_M_ when dA is opposite Fapy•dG (**7b**) is ∼4-fold greater than when a dC-Fapy•dG base pair (**7a**) is extended; this is lower than that measured for a T-Fapy•dG base pair (**7c**). In contrast to when a T-Fapy•dG mispair is extended, *k*_cat_ decreases almost 15-fold when a dA-Fapy•dG mispair is extended compared to reaction of **7a**. The large decrease in extension efficiency for a dA-Fapy•dG mispair (**7b**) compared to a dC-Fapy•dG base pair (**7a**) is different from what is observed for the DNA polymerase I fragment from *E. coli*, Klenow exo^−^ (29). It is also distinct from extension past dA-8-OxodGuo mispairs, which occurs more readily relative to bypass of dC-8-OxodGuo (44,45). In these instances, extension efficiency of the primer past the mispair is far more comparable to that when dC is opposite the lesion, or even greater.

**Table 2.** Extension of primers containing template Fapy•dG by Pol β.^a,b^.

| N | $k_{\text{cat}}$ ($\times 10^{-2} \text{ s}^{-1}$) | $K_M$ ($\mu\text{M}$) | $k_{\text{cat}} / K_M$<br>($\times 10^{-4} \mu\text{M}^{-1} \text{ s}^{-1}$) |
| --- | --- | --- | --- |
| C | $22.4 \pm 2.2$ | $19.3 \pm 3.3$ | $116.0 \pm 23.0$ |
| A | $1.6 \pm 0.2$ | $83.7 \pm 17.1$ | $1.9 \pm 0.5$ |
| T | $31.6 \pm 4.0$ | $138.0 \pm 21.2$ | $22.9 \pm 4.6$ |

<sup>a</sup>[DNA] = 100 nM, [Pol $\beta$ ] = 3 nM. <sup>b</sup>Data are the ave. ± std. dev. of 3 experiments. Each experiment is comprised of 3 replicates.

X-ray crystallography was performed using crystals of Pol β in complex with DNA containing Fapy•dG opposite dC, where dC is in the primer terminal position of the nascent strand. The crystal diffracted to 1.98 Å in space group P2_1_ with Pol β in the open conformation (Figure 5a and Table S1). Fapy•dG base pairs with dC in a Watson-Crick manner with hydrogen bonds between O6, N1, and N2 of Fapy•dG and N4, N3, and O2 of dC, respectively (Figure 5b). The formamide group is in the major groove, uninvolved in base pairing. Ordered water molecules interact with N9 and O4’ of Fapy•dG, and only the β-anomer is observed (Figure 5b). The B-factor of Fapy•dG is 34.7 Å^2^, slightly higher than the B-factor of dC at the primer terminus which is 28.3 Å^2^. Interestingly, the cytosine 3’-OH has poor density and a B-factor of 52.8 Å^2^, substantially higher compared to the rest of the base (B-factor_avg_ = 28.3 Å^2^), which is consistent with reduced extension efficiency.

**Figure 5.**
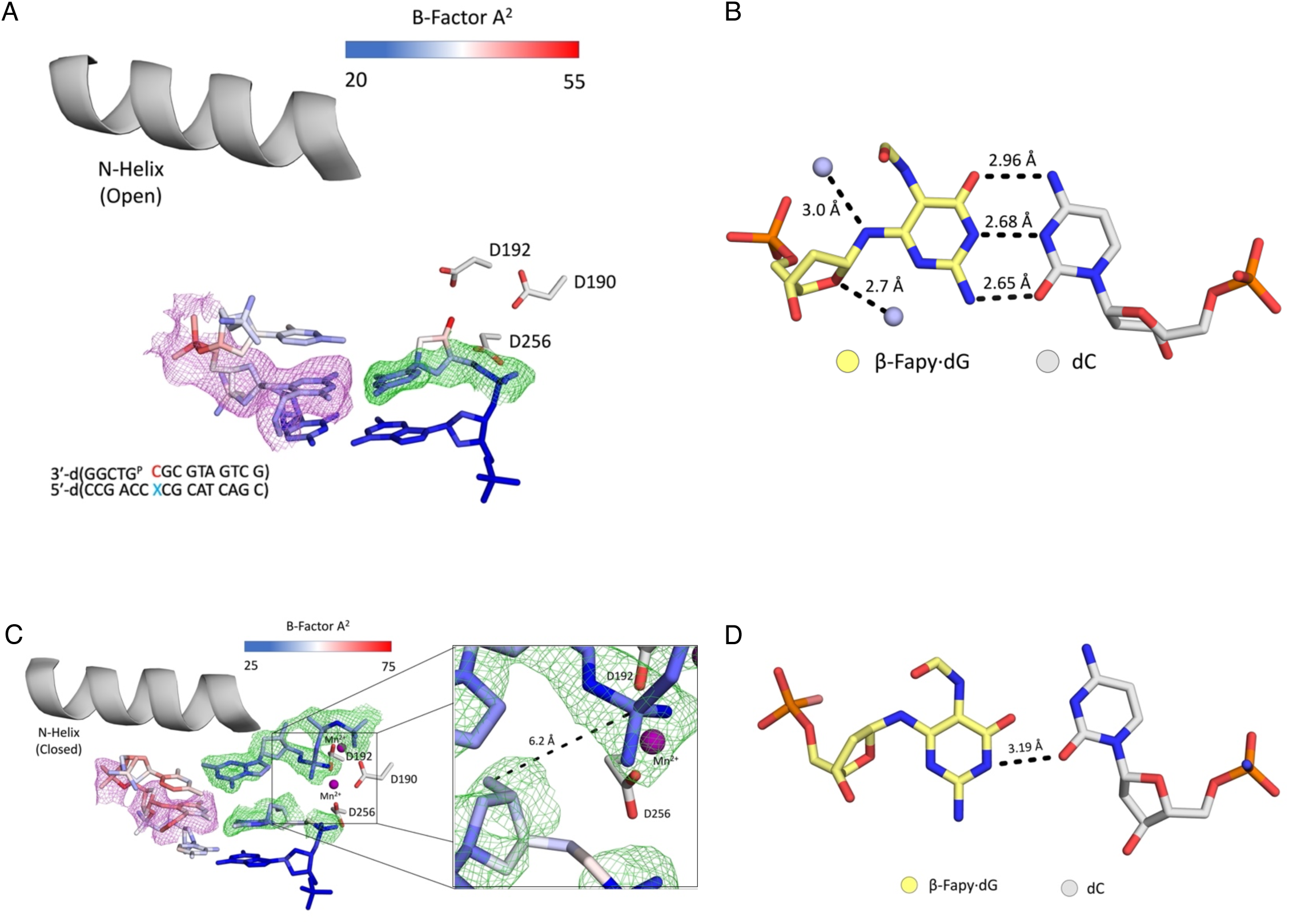
Structural analysis of dC-Fapy•dG extension. **(A)** Active site of the binary Pol β complex. A legend for the B-factors is shown in a blue-white-red gradient. **(B)** β-Fapy•dG (yellow sticks) base paired with dC (gray sticks). Two ordered water molecules are within hydrogen bonding distance of N9 and O4’ of Fapy•dG. **(C)** Ternary Pol β complex with an incoming dGTP. The inset shows a focused view of the 3′-OH and α-PO4. **(D)** β-Fapy•dG (yellow sticks) base paired with dC (gray sticks) from the ternary complex. Manganese ions are colored purple. Omit maps are shown as green mesh contoured to 2σ. Polder maps are shown as purple mesh contoured to 2σ. Key distances are shown as dashes and labeled.

Further characterization of extension was accomplished by solving a ternary structure of dC-Fapy•dG with an incoming nonhydrolyzable dGTP analog (dGpCpp). Binary crystals were soaked in a cryo-solution containing 1 mM dGpCpp and 10 mM MnCl_2_. The resulting crystal diffracted to 2.05 Å in space group P2_1_ (Figure 5c and Table S1). This structure shows Pol β adopts the closed conformation, with the N-helix closed around the incoming dGpCpp. Like the binary structure, Fapy•dG exists in the β-anomer. This Fapy•dG stereoisomer forms a single hydrogen bond with the primer terminal dC, in contrast to the Watson-Crick base pairing observed in the dC-Fapy•dG binary structure (Figure 5d). This altered base pairing leads to substantial conformational changes in the primer terminal dC across from Fapy•dG. This change positions the 3’-OH in the minor groove, 6.2 Å from the α-PO_4_ and does not coordinate the catalytic metal (Figure S6a).Therefore, the overall organization of the active site containing the primer terminal dC when base paired with Fapy•dG provides a strong rationale for the reduction in catalytic efficiency observed in the kinetics (**7a**, Table 2).

Similar structural analysis was performed with Pol β:DNA crystals featuring dA at the primer terminus across from Fapy•dG (dA-Fapy•dG). This binary structure was solved to 1.90 Å in space group P2_1_ with Pol β in the open conformation (Figure 6a and Table S1). Fapy•dG adopts both α- and β-anomers when base paired with dA, with the occupancy of β-Fapy•dG at 31% and α-Fapy•dG at 69% (Figure 6a). Both anomers of Fapy•dG are within hydrogen bond distance of dA. Specifically, the N1 of dA and the N1 of Fapy·dG, as well as the N6 of dA and the O6 of Fapy·dG (Figure 6b). A single ordered water molecule is observed interacting with O4’ and N3 of the Fapy•dG anomers (Figure 6b). Previous analysis suggests that a water in this position can assist in epimerization (27). The formamide group of both α-and β-Fapy•dG is positioned in the major groove and hydrogen bonds with the 3’-flanking dC (Figure 6a). The poor density at the 3’-OH of Fapy·dG indicate the primer terminal base pair is dynamic, which is consistent with the observed inefficient catalysis of primer extension (**7b**, Table 2).

**Figure 6.**
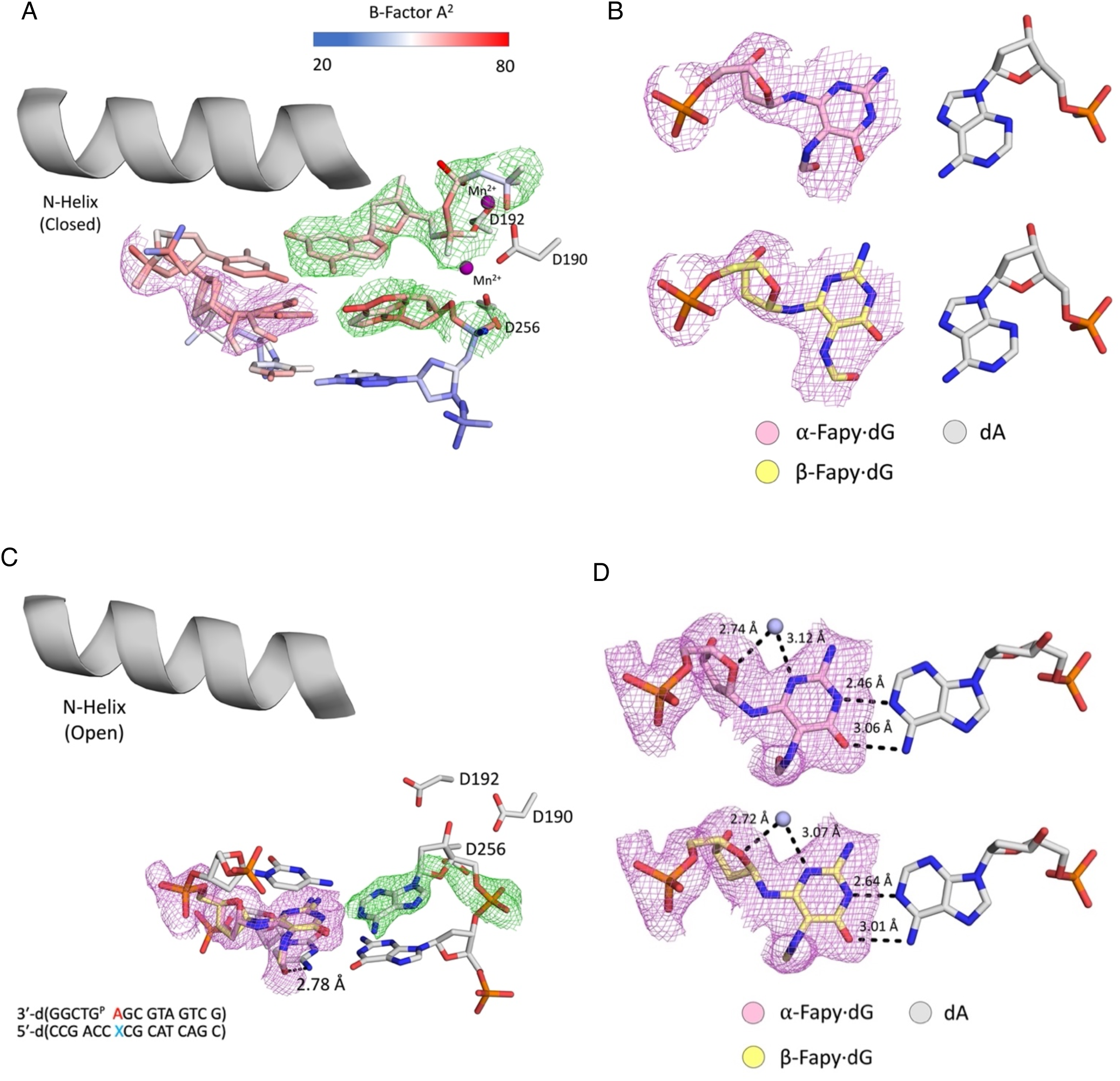
Structural analysis of dA-Fapy•dG extension. **(A)** Active site of the binary Pol β complex. **(B)** α-Fapy•dG (pink sticks) and β-Fapy•dG (yellow sticks) base paired with dA (gray sticks). An ordered water molecule is within hydrogen bonding distance of O4’ and N3 of both Fapy•dG anomers. **(C)** Ternary Pol β complex with an incoming dGpCpp. A legend for the B-factors is depicted in a blue-white-red gradient. **(D)** α-and β-Fapy•dG (pink and yellow sticks, respectively) base paired with dA (gray sticks). Manganese ions are colored purple. Omit maps are shown as green mesh contoured to 2σ. Polder maps are shown as purple mesh contoured to 2σ. Key distances are shown as dashes and labeled.

To better understand the cause of the reduced efficiency for extension from dA-Fapy•dG (Table 2), we solved a ternary structure with incoming nonhydrolyzable dGTP (dGpCpp) to identify perturbations in the active site that may hinder catalysis. To that end, binary crystals were soaked in a cryo-solution containing 1 mM dGpCpp and 10 mM MnCl_2_. The crystal diffracted to 2.64 Å and was solved in space group P2_1_ with Pol β in a closed conformation (Figure 6c). Fapy•dG adopts both α-and β-anomers when base paired with dA, with the occupancy of β-Fapy•dG at 42% and α-Fapy•dG at 58% (Figure 6c). To accommodate binding of the dGpCpp, rotation about the glycosidic bond of the primer terminal dA occurs to present its Hoogsteen face to the opposing Fapy•dG (Figure 6d). This distorts the active site of the primer terminal dA-Fapy•dG base pair (Figure 6c). The primer terminal dA and Fapy•dG are separated in the active site to such an extent that they are effectively no longer within hydrogen bonding distance (Figure 6d). The 3’-OH does not coordinate the catalytic metal, and has shifted 5.9 Å away from the α-PO_4_ of the incoming dGpCpp (Figure S7). The presence of an incoming nucleotide does not stabilize the primer terminal dA, which features a high B-factor of 59.7 Å^2^ compared to the surrounding DNA residues (B-factor_avg_ = 42.0 Å^2^). These conformational changes in the Pol β active site poorly position the reactive entities to carry out catalysis, and are consistent with the inefficient primer extension observed when dA is opposite Fapy•dG (Table 2).

Extension from a T-Fapy•dG base pair, features the highest *K_m_* for an incoming nucleotide (Table 2). We were unable to obtain a ternary crystal structure that would provide mechanistic insight to this phenomenon. Valuable information was gleaned from the corresponding binary structure, which diffracted to 2.48 Å in space group P2_1_ (Figure 7a). Due to lack of a distinct conformation in the density of the omit map for Fapy•dG, the configurational isomer cannot be determined. The lesion is modeled in the β-anomer due to the predominance of that configuration in the Fapy•dG-T host guest complex (Figure 4c). The O2 of T and N2 of Fapy•dG are within hydrogen bonding distance and the formamide group of Fapy•dG is in the major groove of DNA (Figure 7b). The average B-factor of Fapy•dG is 67.2 Å^2^, which is strikingly higher than the average B-factor of T at the primer terminus (B-factor_avg_ = 48.6 Å^2^). This relative instability does not create an amenable binding pocket for an incoming nucleotide.

**Figure 7.**
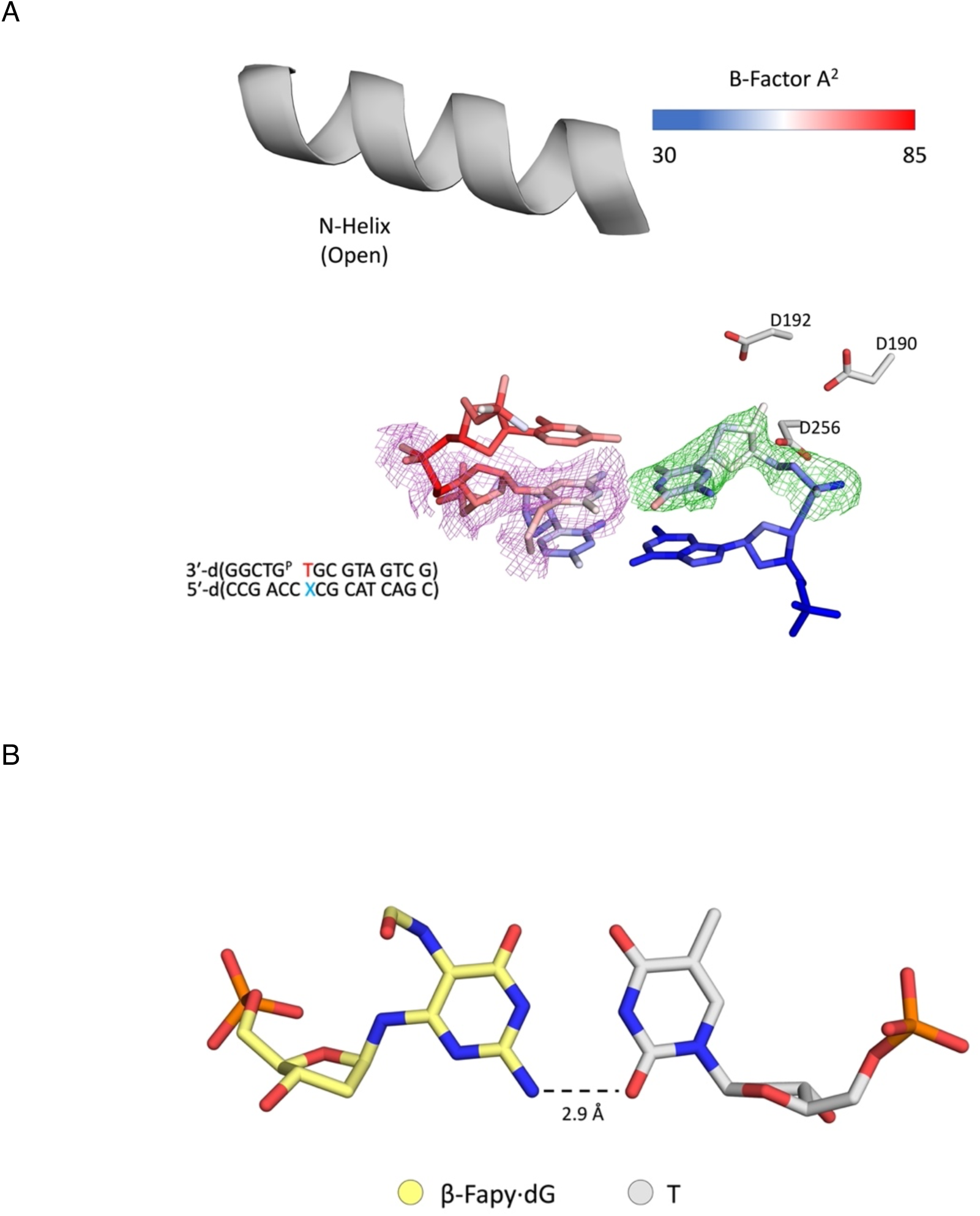
Fapy•dG opposite T at the primer terminus. **(A)** Active site of binary Pol β complex. A legend for the B-factors is shown as a blue-white-red gradient. **(B)** β-Fapy•dG (yellow sticks) opposite the primer terminal T (gray sticks). Omit map is shown as green mesh contoured to 2σ. Polder map is shown in purple mesh contoured to 2σ.

## DISCUSSION

Pol β was used as a model human DNA polymerase to examine replication of Fapy•dG, an important DNA lesion produced from dG oxidation. Complementary biochemical and structural experiments were carried out to examine the role of Fapy•dG as a promutagenic DNA lesion. The possibilities that Fapy•dG could induce mistakes by Pol β via its presence in a DNA template or due to its presence in the nucleotide triphosphate pool were investigated.

Fapy•dGMP incorporation proved to be inefficient, even when base paired non-mutagenically with a template dC. The concentration of dGTP in cells is typically less than 10 μM (43). Hence, even if in the unlikely event Fapy•dGTP concentration approached 10% of this nucleotide, the concentration of the damaged triphosphate would be far below the *K*_M_ values measured (Table 1) for Fapy•dGMP incorporation opposite dC, dA or T. We were unable to obtain structural data for incorporation of natural Fapy•dGTP opposite any templating base. This likely stems from the slow rate of catalysis, and very high *K*_M_ values possibly caused by the gating mechanisms of Pol β’s active site (37).

If Fapy•dGMP was incorporated opposite dA, T, or dC, only when the lesion is paired with the latter, which will not promote mutagenesis, is the complex likely to be extended by Pol β. X-ray crystallography provided a structural rationale for this in that only when Fapy•dG is opposite dC in a template is the 3′-OH in position for catalysis to occur. When a 3′-primer terminal Fapy•dG is part of a promutagenic base pair, a dramatic increase in B-factor for the lesion is observed indicating increased interatomic movement. Increased dynamics of the 3’-hydroxyl group of nascent primers were frequently observed in structures containing Fapy•dG. This environment should make it challenging for the enzyme to efficiently incorporate a nucleotide. Further evidence of this is the inability for Pol β to properly adopt its closed conformation when Fapy•dG-dA is present at the primer terminus. Overall, biochemical and X-ray crystallography experiments suggest that Fapy•dGMP is unlikely to be incorporated via the corresponding nucleotide triphosphate and extended by Pol β and therefore not a source of mutations. Although the activity of MTH1 on Fapy•dGTP is not reported, the triphosphate is a poor substrate for the bacterial analogue (MutT). Inefficient incorporation and extension by Pol β suggests that nature does not need the dNTPase to protect against Fapy•dG mutagenicity (36,46). However, future studies will need to be performed to investigate this.

Replication studies in HEK 293T cells revealed that Fapy•dG bypass results in misincorporation of dA and T (23). Insertion of dCMP and dAMP opposite Fapy•dG in the template position by Pol β has been studied (27). This investigation provided the first insight into thymidine incorporation opposite the lesion. Interestingly, TMP insertion is less efficient than that of dCMP and dAMP. Pol β is unable to properly close when TTP is the incoming nucleotide and the nucleotide triphosphate does not form any hydrogen bonds with the template Fapy•dG. These structural features indicate why TMP incorporation is least efficient and has the highest *K*_M_ value. Once incorporated in duplex DNA, the T-Fapy•dG base pair does not perturb the DNA helix and Fapy•dG adopts both the α-and β-anomers. Fapy•dG behaves similarly when the lesion base pairs with dA in duplex DNA (27).

When an incorporation event occurs opposite Fapy•dG, large reductions in catalytic efficiency compared to undamaged nucleotides are observed for extension from dC, dA, and dT (250-, 15,000-, and, 1,200-fold reductions, respectively) (27). The primary observation that provides a rationale for these kinetic effects stems from dynamics induced by the presence of the lesion at the primer terminus. Extension from the dC-Fapy•dG base pair occurs most efficiently, but even in this nonmutagenic pairing the lesion disrupts positioning for efficient catalysis. Our structural studies reveal that the the 3′-OH does not coordinate the catalytic metal and shifts away from the α-PO_4_, preventing efficient catalysis. A more severe distortion occurs in the dA-β-Fapy•dG base pair where rotation about the dA glycosidic bond results in the presentation of its Hoogsteen face; creating a dynamic primer terminus as evidenced by elevated B-factor of the primer terminal dA. Furthermore, the 3′-OH is no longer in position for catalysis, dramatically reducing the ability to extend from this base pair. For the T-Fapy•dG binary complex we also observe instability at the primer terminus, indicating that further extension is unlikely. Together, extension from a native nucleotide base paired with Fapy•dG is largely hindered due to induced dynamics of the primer terminal base by the opposing lesion. Overall, stereochemical dynamics due to configurational and conformational changes contribute significantly to the effects of Fapy•dG on the polymerase activity of DNA polymerase β. These properties indicate that Fapy•dG bypass by Pol β is significantly more challenging than the less mutagenic related DNA lesion, 8-OxodGuo (45). Additional studies involving bypass polymerases could shed further light on the promutagenic proclivity of Fapy•dG.

## Data availability

The atomic coordinate and structure factor for all the x-ray crystal structurs reported here have been deposited in the Protein Data Bank under Accession Codes 8VFA, 8VFB, 8VFC, 8VFD, 8VFE, 8VFF, 8VFG, 8VFH, 8VFI, 8VFJ, 8VF8, and 8VF9.

## Supplemental data

Supplemental data are available at NAR online

## References

1. Cadet, J., Douki, T. and Ravanat, J.-L. (2008) Oxidatively Generated Damage to the Guanine Moiety of DNA: Mechanistic Aspects and Formation in Cells. Acc. Chem. Res., 41, 1075–1083.

2. Dizdaroglu, M. and Jaruga, P. (2012) Mechanisms of free radical-induced damage to DNA. Free Rad. Res., 46, 382–419.

3. Dizdaroglu, M., Coskun, E. and Jaruga, P. (2017) Repair of oxidatively induced DNA damage by DNA glycosylases: Mechanisms of action, substrate specificities and excision kinetics. Mutat. Rev. Res. Mutagen., 771, 99–127.

4. Delaney, J.C. and Essigmann, J.M. (2008) Biological Properties of Single Chemical-DNA Adducts: A Twenty Year Perspective. Chem. Res. Toxicol., 21, 232–252.

5. Ravanat, J.L., Douki, T., Duez, P., Gremaud, E., Herbert, K., Hofer, T., Lasserre, L., Saint-Pierre, C., Favier, A. and Cadet, J. (2002) Cellular background level of 8-oxo-7,8-dihydro-2’-deoxyguanosine: an isotope based method to evaluate artefactual oxidation of DNA during its extraction and subsequent work-up. Carcinogenesis, 23, 1911–1918.

6. Fleming, A.M. and Burrows, C.J. (2017) 8-Oxo-7,8-dihydroguanine, friend and foe: Epigenetic-like regulator versus initiator of mutagenesis. DNA Repair, 56, 75–83.

7. McAuley-Hecht, K.E., Leonard, G.A., Gibson, N.J., Thomson, J.B., Watson, W.P., Hunter, W.N. and Brown, T. (1994) Crystal Structure of a DNA Duplex Containing 8-Hydroxydeoxyguanine-Adenine Base Pairs. Biochemistry, 33, 10266–10270.

8. Evans, M.D., Dizdaroglu, M. and Cooke, M.S. (2004) Oxidative DNA damage and disease: induction, repair and significance. Mutat. Res., 567, 1–61.

9. Fleming, A.M., Ding, Y. and Burrows, C.J. (2017) Oxidative DNA damage is epigenetic by regulating gene transcription via base excision repair. Proc. Natl. Acad. Sci. USA, 114, 2604–2609.

10. Wu, J., McKeague, M. and Sturla, S.J. (2018) Nucleotide-Resolution Genome-Wide Mapping of Oxidative DNA Damage by Click-Code-Seq. J. Am. Chem. Soc., 140, 9783–9787.

11. Graille, M., Wild, P., Sauvain, J.-J., Hemmendinger, M., Guseva Canu, I. and Hopf, N.B. (2020) Urinary 8-OHdG as a Biomarker for Oxidative Stress: A Systematic Literature Review and Meta-Analysis. Int. J. Mol. Sci., 21, 3743.

12. Tan, X., Grollman, A.P. and Shibutani, S. (1999) Comparison of the Mutagenic Properties of 8-Oxo-7,8-dihydro-2’-deoxyadenosine and 8-Oxo-7-8-dihydro-2’-deoxyguanosine DNA Lesions in Mammalian Cells. Carcinogenesis, 20, 2287–2292.

13. Greenberg, M.M. (2012) The Formamidopyrimidines: Purine Lesions Formed in Competition With 8-Oxopurines From Oxidative Stress. Acc. Chem. Res., 45, 588–597.

14. Douki, T., Martini, R., Ravanat, J.-L., Turesky, R.J. and Cadet, J. (1997) Measurement of 2,6-Diamino-4-hydroxy-5-formamidopyrimidine and 8-Oxo-7,8-dihydroguanine in Isolated DNA Exposed to Gamma Radiation in Aqueous Solution. Carcinogenesis, 18, 2385–2391.

15. Jaruga, P., Kirkali, G. and Dizdaroglu, M. (2008) Measurement of Formamidopyrimidines in DNA. Free Rad. Biol. Med., 45, 1601–1609.

16. Dizdaroglu, M., Kirkali, G. and Jaruga, P. (2008) Formamidopyrimidines in DNA: Mechanisms of Formation, Repair, and Biological Effects. Free Rad. Biol. Med., 45, 1610–1621.

17. Pouget, J.-P., Douki, T., Richard, M.-J. and Cadet, J. (2000) DNA Damage Induced in Cells by g and UVA Radiation As Measured by HPLC/GC-MS and HPLC-EC and Comet Assay. Chem. Res. Toxicol., 13, 541–549.

18. Liu, D., Croteau, D.L., Souza-Pinto, N., Pitta, M., Tian, J., Wu, C., Jiang, H., Mustafa, K., Keijzers, G., Bohr, V.A. et al. (2010) Evidence that OGG1 Glycosylase Protects Neurons against Oxidative DNA Damage and Cell Death under Ischemic Conditions. J. Cereb. Blood Flow Metab., 31, 680–692.

19. Arczewska, K.D., Tomazella, G.G., Lindvall, J.M., Kassahun, H., Maglioni, S., Torgovnick, A., Henriksson, J., Matilainen, O., Marquis, B.J., Nelson, B.C. et al. (2013) Active transcriptomic and proteomic reprogramming in the C. elegans nucleotide excision repair mutant xpa-1. Nucleic Acids Res., 41, 5368–5381.

20. Tomar, R., Minko, I.G., Sharma, P., Kellum, A.H., Jr., Lei, L., Harp, Joel M., Iverson, T.M., Lloyd, R S., Egli, M. and Stone, Michael P. (2023) Base excision repair of the N-(2-deoxy-d-erythro-pentofuranosyl)-urea lesion by the hNEIL1 glycosylase. Nucleic Acids Res., 51, 3754–3769.

21. Yang, H., Tang, J.A. and Greenberg, M.M. (2020) Synthesis of Oligonucleotides Containing the N6-(2-Deoxy-α,β-D-erythropentofuranosyl)-2,6-diamino-4-hydroxy-5-formamidopyrimidine (Fapy.dG) Oxidative Damage Product Derived from 2’-Deoxyguanosine. Chem. -Eur. J., 26, 5441–5448.

22. Haraguchi, K. and Greenberg, M.M. (2001) Synthesis of Oligonucleotides Containing Fapy•dG (N6-(2-Deoxy-a,b-D-erythro-pento-furanosyl)-2,6-diamino-4-hydroxy-5-formamidopyrimidine). J. Am. Chem. Soc., 123, 8636–8637.

23. Bacurio, J.H.T., Yang, H., Naldiga, S., Powell, B.V., Ryan, B.J., Freudenthal, B.D., Greenberg, M.M. and Basu, A.K. (2021) Sequence context effects of replication of Fapy•dG in three mutational hot spot sequences of the p53 gene in human cells. DNA Repair, 108, 103213.

24. Kalam, M.A., Haraguchi, K., Chandani, S., Loechler, E.L., Moriya, M., Greenberg, M.M. and Basu, A.K. (2006) Genetic Effects of Oxidative DNA Damages: Comparative Mutagenesis of the Imidazole Ring-Opened Formamidopyrimidines (Fapy Lesions) and 8-Oxo-purines in Simian Kidney Cells. Nucleic Acids Res., 34, 2305–2315.

25. Stanio, S., Bacurio, J.H.T., Yang, H., Greenberg, M.M. and Basu, A.K. (2023) 8-Oxo-2′-deoxyguanosine Replication in Mutational Hot Spot Sequences of the p53 Gene in Human Cells Is Less Mutagenic than That of the Corresponding Formamidopyrimidine. Chem. Res. Toxicol., 36, 782–789.

26. Neeley, W.L. and Essigmann, J.M. (2006) Mechanisms of Formation, Genotoxicity, and Mutation of Guanine Oxidation Products. Chem. Res. Toxicol., 19, 491–505.

27. Ryan, B.J., Yang, H., Bacurio, J.H.T., Smith, M.R., Basu, A.K., Greenberg, M.M. and Freudenthal, B.D. (2022) Structural Dynamics of a Common Mutagenic Oxidative DNA Lesion in Duplex DNA and during DNA Replication. J. Am. Chem. Soc., 144, 8054–8065.

28. Gehrke, T.H., Lischke, U., Gasteiger, K.L., Schneider, S., Arnold, S., Müller, H.C., Stephenson, D.S., Zipse, H. and Carell, T. (2013) Unexpected non-Hoogsteen-based mutagenicity mechanism of FaPy-DNA lesions. Nature Chem. Biol., 9, 455–461.

29. Wiederholt, C.J. and Greenberg, M.M. (2002) Fapy•dG Instructs Klenow Exo^−^ to Misincorporate Deoxyadenosine. J. Am. Chem. Soc., 124, 7278–7279.

30. Jun, Y.W., Kant, M., Coskun, E., Kato, T.A., Jaruga, P., Palafox, E., Dizdaroglu, M. and Kool, E.T. (2023) Possible Genetic Risks from Heat-Damaged DNA in Food. ACS Cent. Sci., 9, 1170–1179.

31. Freudenthal, B.D., Beard, W.A., Perera, L., Shock, D.D., Kim, T., Schlick, T. and Wilson, S.H. (2015) Uncovering the polymerase-induced cytotoxicity of an oxidized nucleotide. Nature, 517, 635–639.

32. Whitaker, A.M., Smith, M.R., Schaich, M.A. and Freudenthal, B.D. (2017) Capturing a mammalian DNA polymerase extending from an oxidized nucleotide. Nucleic Acids Res., 45, 6934–6944.

33. Gad, H., Koolmeister, T., Jemth, A.-S., Eshtad, S., Jacques, S.A., Strom, C.E., Svensson, L.M., Schultz, N., Lundback, T., Einarsdottir, B.O. et al. (2014) MTH1 inhibition eradicates cancer by preventing sanitation of the dNTP pool. Nature, 508, 215–221.

34. Huber, K.V.M., Salah, E., Radic, B., Gridling, M., Elkins, J.M., Stukalov, A., Jemth, A.-S., Gokturk, C., Sanjiv, K., Stromberg, K. et al. (2014) Stereospecific targeting of MTH1 by (S)-crizotinib as an anticancer strategy. Nature, 508, 222–227.

35. Lee, Y., Onishi, Y., McPherson, L., Kietrys, A.M., Hebenbrock, M., Jun, Y.W., Das, I., Adimoolam, S., Ji, D., Mohsen, M.G. et al. (2022) Enhancing Repair of Oxidative DNA Damage with Small-Molecule Activators of MTH1. ACS Chem. Biol., 17, 2074–2087.

36. Imoto, S., Patro, J.N., Jiang, Y.L., Oka, N. and Greenberg, M.M. (2006) Synthesis, DNA Polymerase Incorporation, and Enzymatic Phosphate Hydrolysis of Formamidopyrimidine Nucleoside Triphosphates. J. Am. Chem. Soc., 128, 14606–14611.

37. Smith, M.R., Shock, D.D., Beard, W.A., Greenberg, M.M., Freudenthal, B.D. and Wilson, S.H. (2019) A guardian residue hinders insertion of a Fapy•dGTP analog by modulating the open-closed DNA polymerase transition. Nucleic Acids Res., 47, 3197–3207.

38. Scaringe, S.A. (2000) Advanced 5’-silyl-2’-orthoester approach to RNA oligonucleotide synthesis. Methods Enzymol., 317, 3–18.

39. Beard, W.A.W., S. H. (1995) Purification and domain-mapping of mammalian DNA polymerase beta. Methods in enzymology, 262, 98–107.

40. Creighton, S., Bloom, L.B. and Goodman, M.F. (1995) Gel Fidelity Assay Measuring Nucleotide Misinsertion, Exonucleolytic Proofreading, and Lesion Bypass Efficiencies. Methods Enzymol., 262, 232–256.

41. Beard, W.A., Shock, D.D. and Wilson, S.H. (2004) Influence of DNA structure on DNA polymerase beta active site function: extension of mutagenic DNA intermediates. J. Biol. Chem., 279, 31921–31929.

42. Brown, J.A., Duym, W.W., Fowler, J.D. and Suo, Z. (2007) Single-turnover Kinetic Analysis of the Mutagenic Potential of 8-Oxo-7,8-dihydro-2′-deoxyguanosine during Gap-filling Synthesis Catalyzed by Human DNA Polymerases λ and β. J. Mol. Biol., 367, 1258–1269.

43. Traut, T.W. (1994) Physiological concentrations of purines and pyrimidines. Molecular & Cellular Biochemistry, 140, 1–22.

44. Shibutani, S., Takeshita, M. and Grollman, A.P. (1991) Insertion of Specific Bases During DNA Synthesis Past the Oxidation-Damaged Base 8-OxodG. Nature, 349, 431–434.

45. Reed, A.J. and Suo, Z. (2017) Time-Dependent Extension from an 8-Oxoguanine Lesion by Human DNA Polymerase Beta. J. Am. Chem. Soc., 139, 9684–9690.

46. Maki, H. and Sekiguchi, M. (1992) MutT Protein Specifically Hydrolyzes a Potent Mutagenic Substrate for DNA Synthesis. Nature, 355, 273–275.

