## Supplemental information for "Biochemical and Structural Characterization of Fapy•dG Replication by Human DNA Polymerase β"

**Supporting Information: Biochemical and Structural Characterization of the Replication of the Deoxyguanosine Derived Formamidopyrimidine Lesion by Human DNA Polymerase β**

Shijun Gao^a,#^, Peyton N. Oden^b,#^, Benjamin J. Ryan^b^, Haozhe Yang^a,c^, Bret D. Freudenthal^*b^ and Marc M. Greenberg*^a^

^a^Department of Chemistry, Johns Hopkins University, 3400 N. Charles St., Baltimore, MD 21218, USA

^b^Department of Biochemistry and Molecular Biology, and Department of Cancer Biology, University of Kansas Medical Center, Kansas City, Kansas 66160, USA

^c^Current address: Frontiers Medical Center, Tianfu Jincheng Laboratory, Chengdu, Sichuan 610212, China

#These authors contributed equally to this manuscript.

^*^Corresponding author: Bret D. Freudenthal

^*^Corresponding author: Marc M. Greenberg

A

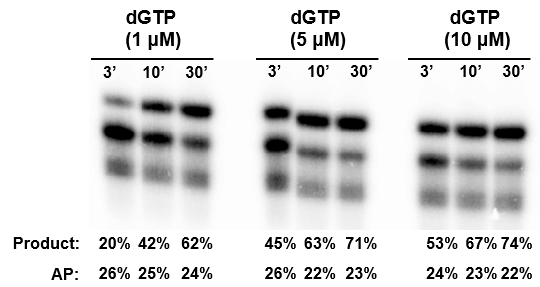

B

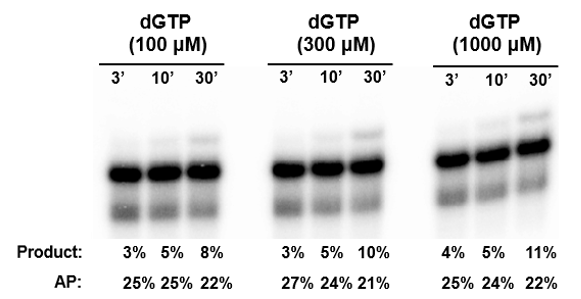

C

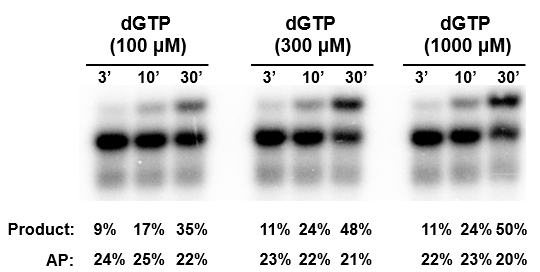

**Figure S1.** **Pol β mediated time-course extension.** (A) Extension of ternary DNA complex containing Fapy·dG opposite dC (**4a**). (B) Extension of ternary DNA complex containing Fapy·dG opposite dA (**4b**). (C) Extension of ternary DNA complex containing Fapy·dG opposite dT (**4c**). Unchanged level of abasic site (AP, lower band) suggested that the reaction was unaffected by the impurity in the primer.

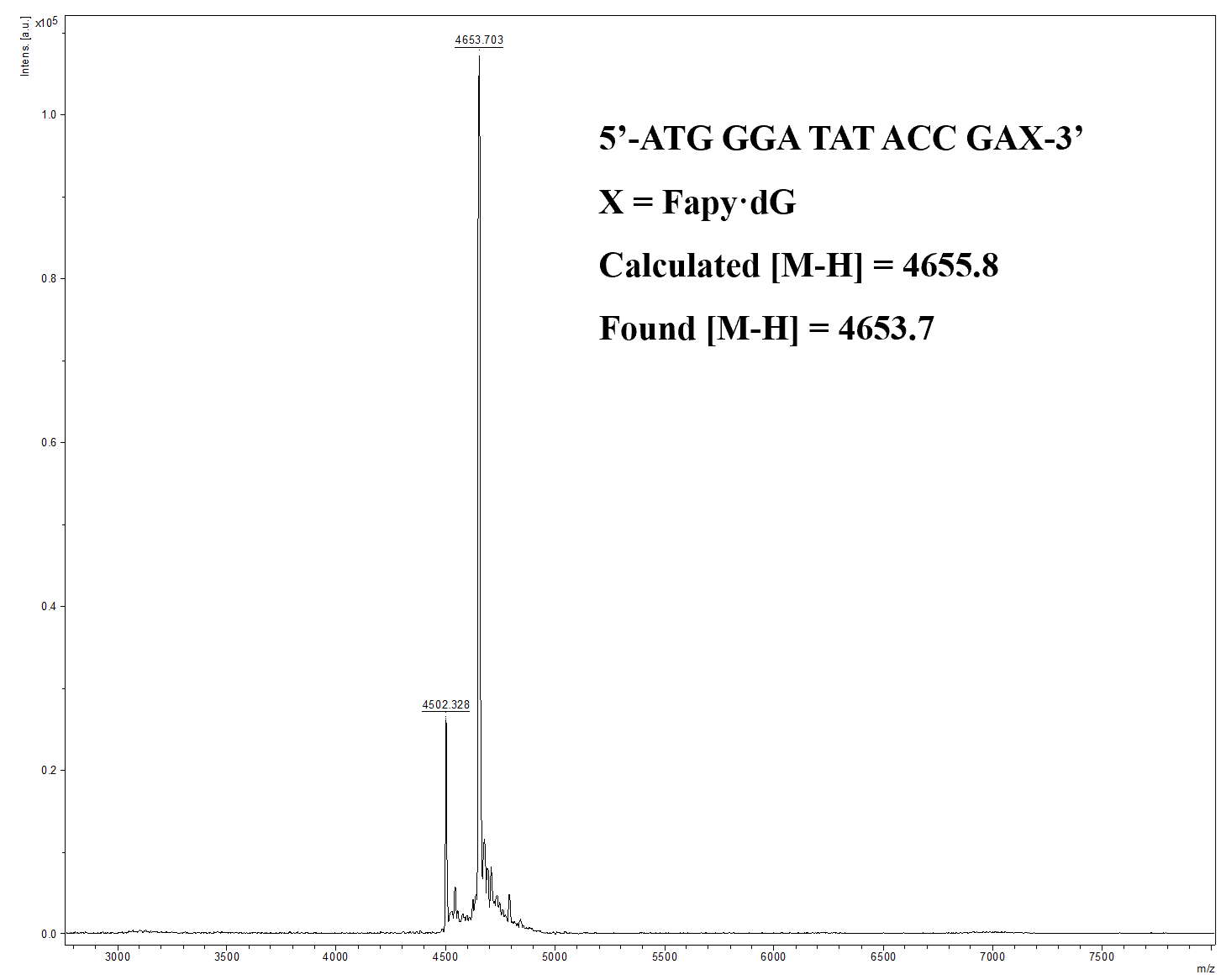

**Figure S2**. MALDI-TOF MS of 3’-Fapy·dG-containing primer in **4a**-**c**.

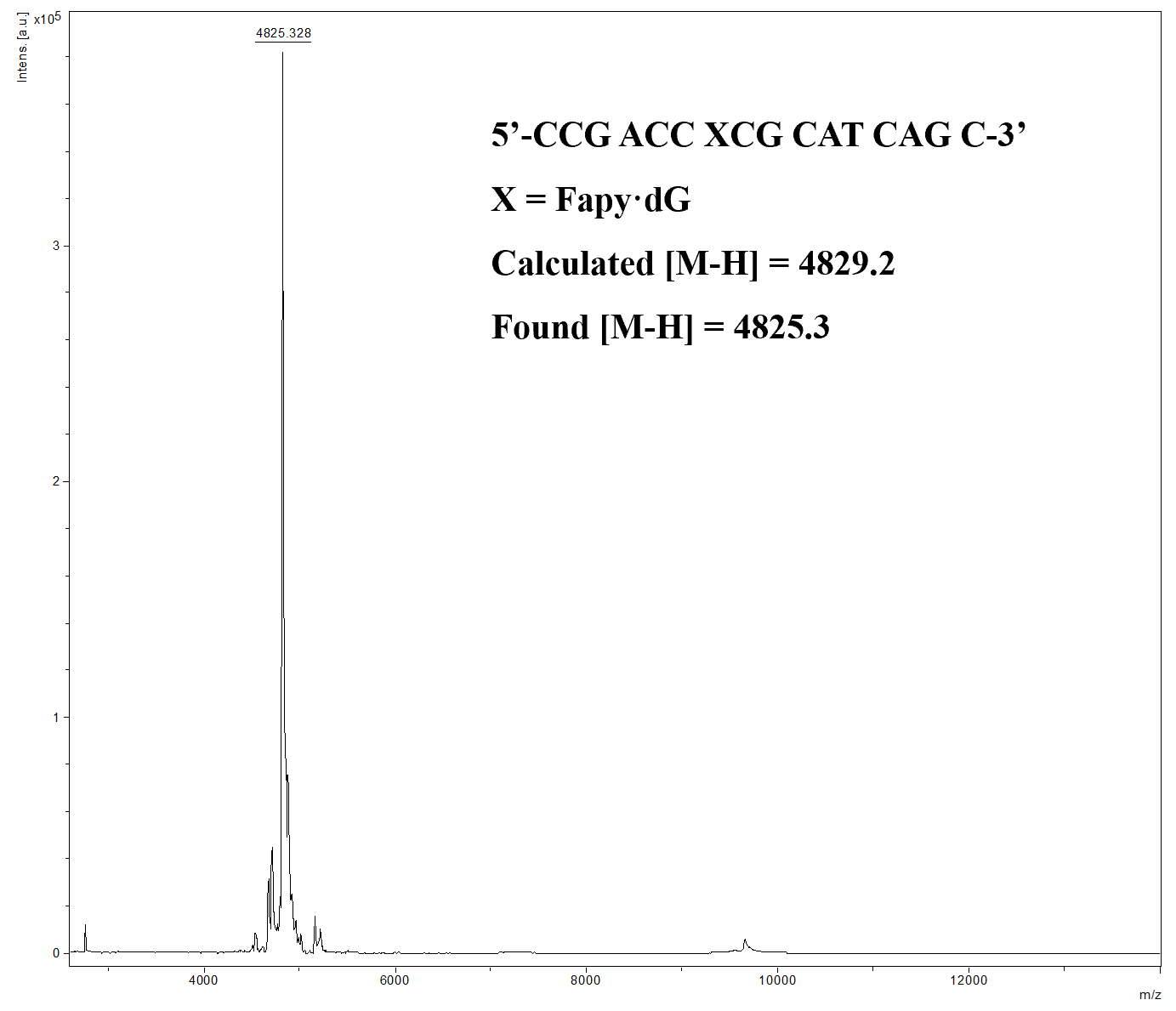

**Figure S3**. MALDI-TOF MS of template oligonucleotide with Fapy·dG opposite the primer terminus used in crystallization.

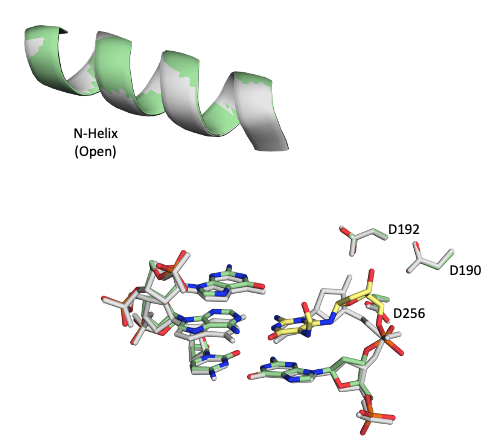

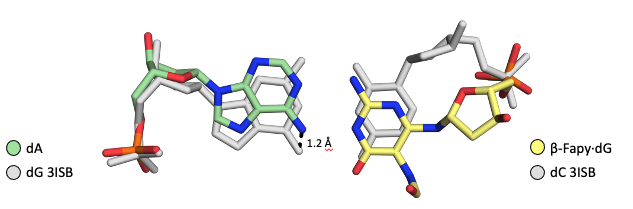

A

B

**Figure S4.** (A) Overlay of dA-Fapy•dG (green and yellow sticks) and a binary Pol β:DNA complex (PDB: 3ISB). (B) Close-up view of overlay between Fapy•dG-dA and 3ISB (gray sticks).

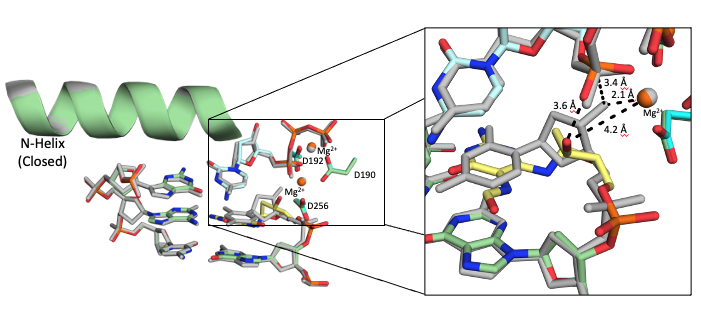

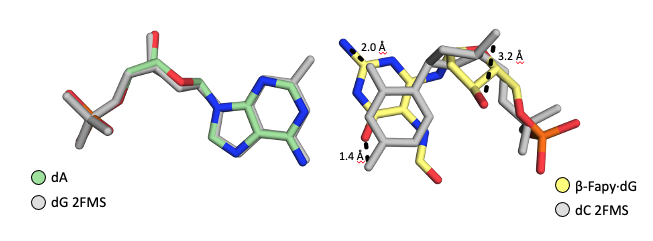

A

B

**Figure S5.** (A) Overlay of dA-Fapy•dG with an incoming dCTP (green, yellow, and cyan sticks) and a ternary Pol β:DNA complex (PDB: 2FMS). Magnesium ions in the catalytic triad for dA-Fapy•dG indicated in orange. Magnesium ions in the catalytic triad for 2FMS indicated in gray. (B) Close-up view of overlay between Fapy•dG-dA and 2FMS (gray sticks).

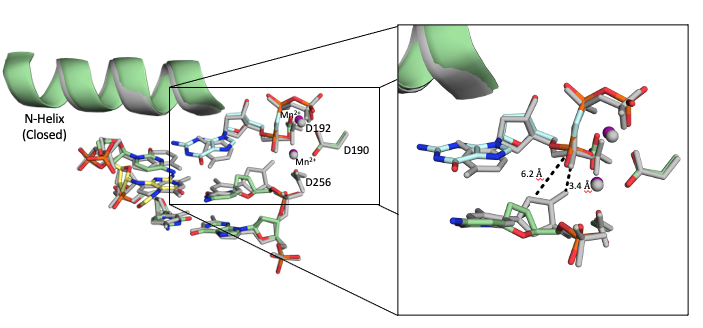

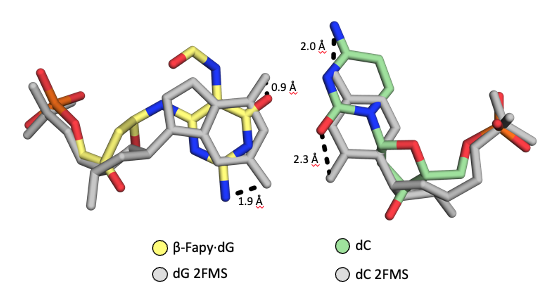

A

B

**Figure S6.** (A) Overlay of dC-Fapy•dG (green and yellow sticks) and a ternary Pol β:DNA complex (PDB: 2FMS). Manganese ions in the catalytic triad for dA-Fapy•dG indicated in purple. Magnesium ions in the catalytic triad for 2FMS indicated in gray.

(B) Close-up view of overlay between dC-Fapy•dG and 2FMS (grey sticks).

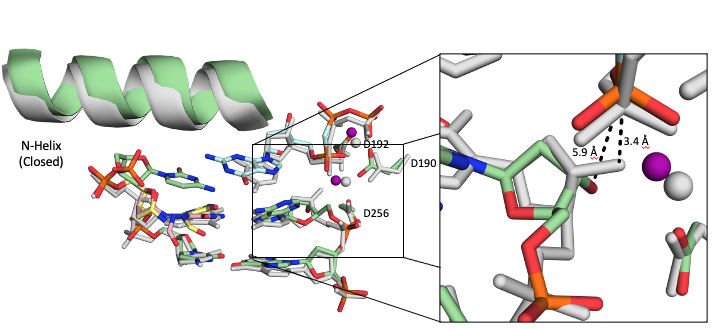

**Figure S7.** Overlay of dA-Fapy•dG (green, pink, and yellow sticks) and a ternary Pol β:DNA complex (PDB: 2FMS). Manganese ions in the catalytic triad for dA-Fapy•dG indicated in purple. Magnesium ions in the catalytic triad for 2FMS indicated in gray.

|  | Pol β Binary  dC-Fapy∙dG | Pol β Binary  dA-Fapy∙dG | Pol β Binary  dT-Fapy∙dG | Pol β Ternary  dC-Fapy∙dG  dGTP |
| --- | --- | --- | --- | --- |
| **Data collection** |  |  |  |  |
| Space group | P2_1_ | P2_1_ | P2_1_ | P2_1_ |
| Cell dimensions |  |  |  |  |
| *a*, *b*, *c* (Å) | 54.13, 79.44, 55.02 | 52.44, 78.63, 54.52 | 54.06, 79.73, 55.68 | 54.44, 79.53, 54.89 |
| *α*, *β*, *γ* (°) | 90, 105.37, 90 | 90, 105.89, 90 | 90, 105.67, 90 | 90, 108.26, 90 |
| Resolution (Å) | 25.00 - 1.98 | 25.00 – 1.90 | 25.00 - 2.48 | 25.00 – 2.05 |
| *R*_meas_^a^ (%) | 0.08 (0.86) | 0.09 (0.95) | 0.08 (0.53) | 0.05 (0.38) |
| *CC_1/2_* ^a^ | (0.47) | (0.49) | (0.85) | (0.83) |
| *I*/σ*I* ^a^ | 12.6 (0.9) | 17.5 (1.3) | 15.1 (2.4) | 24.7 (2.5) |
| Completeness ^a^ (%) | 95.0 | 99.8 | 90.9 | 99.1 |
| Redundancy ^a^ | 2.6 (1.7) | 4.6 (2.7) | 3.4 (3.0) | 4.6 (2.0) |
| **Refinement** |  |  |  |  |
| Resolution (Å) | 1.98 | 1.90 | 2.48 | 2.05 |
| No. reflections | 32838 | 30987 | 14431 | 38424 |
| *R*_work_ / *R*_free_ | 0.19 / 0.22 | 0.21 / 0.23 | 0.20 / 0.26 | 0.23 / 0.27 |
| No. atoms | 3508 | 3503 | 3323 | 3337 |
| Protein | 2566 | 2586 | 2589 | 2581 |
| DNA | 607 | 609 | 608 | 607 |
| Water | 308 | 258 | 99 | 91 |
| B-factors (Å^2^) |  |  |  |  |
| Protein | 30.5 | 25.2 | 40.9 | 35.0 |
| DNA | 29.1 | 24.5 | 38.9 | 29.8 |
| Water | 33.9 | 27.9 | 36.1 | 30.1 |
| R.m.s deviations |  |  |  |  |
| Bond length (Å) | 0.01 | 0.01 | 0.01 | 0.01 |
| Bond angles (º) | 1.23 | 1.35 | 1.14 | 1.42 |
| Ramachandran plot: |  |  |  |  |
| Outliers (%) | 0.00 | 0.00 | 0.00 | 0.00 |
| Allowed (%) | 3.72 | 3.42 | 3.41 | 3.16 |
| Favored (%) | 96.28 | 96.58 | 96.59 | 96.84 |
| PDB ID | 8VF8 | 8VF9 | 8VFC | 8VFA |

^a^ Highest resolution shell is shown in parentheses

**Table S1.** X-ray table

|  | Pol β Ternary  dA-Fapy∙dG  dGTP | Pol β Ternary  dTTP-Fapy∙dG | Pol β Binary  Fapy∙dG-dT | Host Guest  Fapy∙dG-dT |
| --- | --- | --- | --- | --- |
| **Data collection** |  |  |  |  |
| Space group | P2_1_ | P2_1_ | P2_1_ | P2_1_ |
| Cell dimensions |  |  |  |  |
| *a*, *b*, *c* (Å) | 53.85, 79.31, 54.70 | 53.77, 79.07, 54.37 | 53.68, 79.29, 54.96 | 55.38, 80.89, 58.30 |
| *α*, *β*, *γ* (°) | 90, 108.88, 90 | 90, 107.91, 90 | 90, 106.15, 90 | 90, 110.15, 90 |
| Resolution (Å) | 25.00 – 2.64 | 25.00 – 2.10 | 25.00 - 2.09 | 25.00 – 2.14 |
| *R*_meas_^a^ (%) | 0.47 (0.00) | 0.07 (0.99) | 0.07 (0.68) | 0.07 (0.67) |
| *CC_1/2_* ^a^ | (0.59) | (0.43) | (0.63) | (0.69) |
| *I*/σ*I* ^a^ | 3.1 (0.4) | 18.9 (1.3) | 16.8 (1.5) | 18.2 (2.2) |
| Completeness ^a^ (%) | 100.0 | 99.5 | 99.9 | 98.5 |
| Redundancy ^a^ | 4.3 (3.9) | 4.0 (2.4) | 4.3 (2.2) | 3.3 (3.1) |
| **Refinement** |  |  |  |  |
| Resolution (Å) | 2.64 | 2.10 | 2.09 | 2.14 |
| No. reflections | 8172 | 32606 | 36973 | 34645 |
| *R*_work_ / *R*_free_ | 0.20 / 0.20 | 0.20 / 0.24 | 0.20 / 0.24 | 0.21 / 0.26 |
| No. atoms | 3263 | 3312 | 3391 | 3397 |
| Protein | 2573 | 2539 | 2593 | 2593 |
| DNA | 609 | 639 | 611 | 611 |
| Water | 81 | 102 | 160 | 144 |
| B-factors (Å^2^) |  |  |  |  |
| Protein | 45.2 | 32.4 | 32.1 | 33.4 |
| DNA | 42.0 | 35.5 | 33.3 | 33.8 |
| Water | 57.7 | 32.6 | 30.9 | 32.5 |
| R.m.s deviations |  |  |  |  |
| Bond length (Å) | 0.01 | 0.01 | 0.01 | 0.01 |
| Bond angles (º) | 1.58 | 1.10 | 1.26 | 1.15 |
| Ramachandran plot: |  |  |  |  |
| Outliers (%) | 0.00 | 0.00 | 0.00 | 0.00 |
| Allowed (%) | 6.96 | 4.10 | 3.41 | 4.33 |
| Favored (%) | 93.04 | 95.90 | 96.59 | 95.67 |
| PDB ID | 8VFB | 8VFD | 8VFE | 8VFJ |

^a^ Highest resolution shell is shown in parentheses

**Table S1 cont.** X-ray table

|  | Pol β Binary  Fapy∙dG-dA | Pol β Binary  Fapy∙dG-dC | Pol β Ternary  Fapy∙dG-dA  dCTP | Pol β Ternary  Fapy∙dG-dC  dCTP |
| --- | --- | --- | --- | --- |
| **Data collection** |  |  |  |  |
| Space group | P2_1_ | P2_1_ | P2_1_ | P2_1_ |
| Cell dimensions |  |  |  |  |
| *a*, *b*, *c* (Å) | 54.40, 79.28, 54.88 | 54.38, 79.24, 54.88 | 50.80, 80.40, 55.50 | 50.80, 80.40, 55.50 |
| *α*, *β*, *γ* (°) | 90, 105.53, 90 | 90, 105.49, 90 | 90, 107.90, 90 | 90, 107.90, 90 |
| Resolution (Å) | 25.00 – 1.69 | 50.00 – 1.54 | 25.00 – 1.77 | 25.00 – 2.01 |
| *R*_meas_^a^ (%) | 0.05 (0.00) | 0.04 (0.00) | 0.08 (0.00) | 0.11 (0.00) |
| *CC_1/2_* ^a^ | (0.43) | (0.38) | (0.42) | (0.40) |
| *I*/σ*I* ^a^ | 17.5 (0.9) | 27.1 (0.9) | 14.4 (0.8) | 11.2 (0.9) |
| Completeness ^a^ (%) | 93.4 | 97.6 | 99.5 | 99.6 |
| Redundancy ^a^ | 1.8 (1.4) | 3.0 (1.8) | 3.8 (2.7) | 4.1 (2.6) |
| **Refinement** |  |  |  |  |
| Resolution (Å) | 1.69 | 1.54 | 1.77 | 2.01 |
| No. reflections | 43590 | 58170 | 37643 | 24631 |
| *R*_work_ / *R*_free_ | 0.21 / 0.23 | 0.20 / 0.24 | 0.18 / 0.23 | 0.18 / 0.25 |
| No. atoms | 3594 | 3827 | 3770 | 3604 |
| Protein | 2589 | 2588 | 2676 | 2658 |
| DNA | 612 | 610 | 612 | 610 |
| Water | 366 | 606 | 423 | 277 |
| B-factors (Å^2^) |  |  |  |  |
| Protein | 21.0 | 19.2 | 23.7 | 29.6 |
| DNA | 22.3 | 20.7 | 33.1 | 40.6 |
| Water | 25.5 | 27.4 | 33.5 | 35.3 |
| R.m.s deviations |  |  |  |  |
| Bond length (Å) | 0.01 | 0.01 | 0.01 | 0.01 |
| Bond angles (º) | 1.25 | 1.04 | 1.29 | 1.04 |
| Ramachandran plot: |  |  |  |  |
| Outliers (%) | 0.00 | 0.00 | 0.00 | 0.00 |
| Allowed (%) | 1.86 | 2.48 | 1.54 | 1.85 |
| Favored (%) | 98.14 | 97.52 | 98.46 | 98.15 |
| PDB ID | 8VFF | 8VFG | 8VFI | 8VFH |

^a^ Highest resolution shell is shown in parentheses

**Table S1 cont.** X-ray table
